# isomiRs-specific differential expression is the rule, not the exception: Are we missing hundreds of species in microRNA analysis?

**DOI:** 10.1101/2021.12.15.472814

**Authors:** Eloi Schmauch, Pia Laitinen, Tiia A. Turunen, Mari-Anna Väänänen, Tarja Malm, Manolis Kellis, Minna U Kaikkonen, Suvi Linna-Kuosmanen

## Abstract

MicroRNAs (miRNAs) are small RNA molecules that act as regulators of gene expression through targeted mRNA degradation. They are involved in many biological and pathophysiological processes and are widely studied as potential biomarkers and therapeutics agents for human diseases, including cardiovascular disorders. Recently discovered isoforms of miRNAs (isomiRs) exist in high quantities and are very diverse. Despite having few differences with their corresponding reference miRNAs, they display specific functions and expression profiles, across tissues and conditions. However, they are still overlooked and understudied, as we lack a comprehensive view on their condition-specific regulation and impact on differential expression analysis. Here, we show that isomiRs can have major effects on differential expression analysis results, as their expression is independent of their host miRNA genes or reference sequences. We present two miRNA-seq datasets from human umbilical vein endothelial cells, and assess isomiR expression in response to senescence and compartment-specificity (nuclear/cytosolic) under hypoxia. We compare three different methods for miRNA analysis, including isomiR-specific analysis, and show that ignoring isomiRs induces major biases in differential expression. Moreover, isomiR analysis permits higher resolution of complex signal dissection, such as the impact of hypoxia on compartment localization, and differential isomiR type enrichments between compartments. Finally, we show important distribution differences across conditions, independently of global miRNA expression signals. Our results raise concerns over the quasi exclusive use of miRNA reference sequences in miRNA-seq processing and experimental assays. We hope that our work will guide future isomiR expression studies, which will correct some biases introduced by golden standard analysis, improving the resolution of such assays and the biological significance of their downstream studies.

## INTRODUCTION

MicroRNAs (miRNAs) are endogenous non-coding RNAs. Their primary role is to target mRNA for translational inhibition or degradation (1). miRNAs are transcribed from the genome by the RNA polymerase II, then cleaved and processed to yield mature miRNA sequences (**Fig 1A**) (2). With the advent of next generation sequencing, and its generalization in miRNA studies, sequences that vary from the established reference miRNAs have been discovered (3). Most of these isoforms, called isomiRs, have modifications in only one or two nucleotides compared to their reference sequences. isomiRs are generated through various mechanisms, such as alternative cleavage by Drosha/Dicer and non-templated nucleotide additions (4–6). These alternative processing mechanisms give rise to different types of modifications that can be used to categorise isomiRs (**Fig 1B**) (5). As isomiRs are an abundant, highly expressed (6–8), and very diverse group of RNA species that display distinct expression patterns between individuals, tissues (9), cell types (10), sex (11), age (12), and diseases (13–16) they are considered as potential biomarkers in various disoders (17–19) as well as potential therapeutic targets and agents (20, 21). However, isomiRs are still marginalized and their inclusion in miRNA studies is anecdotal, although many tools exist for their analysis.

**Figure 1:**
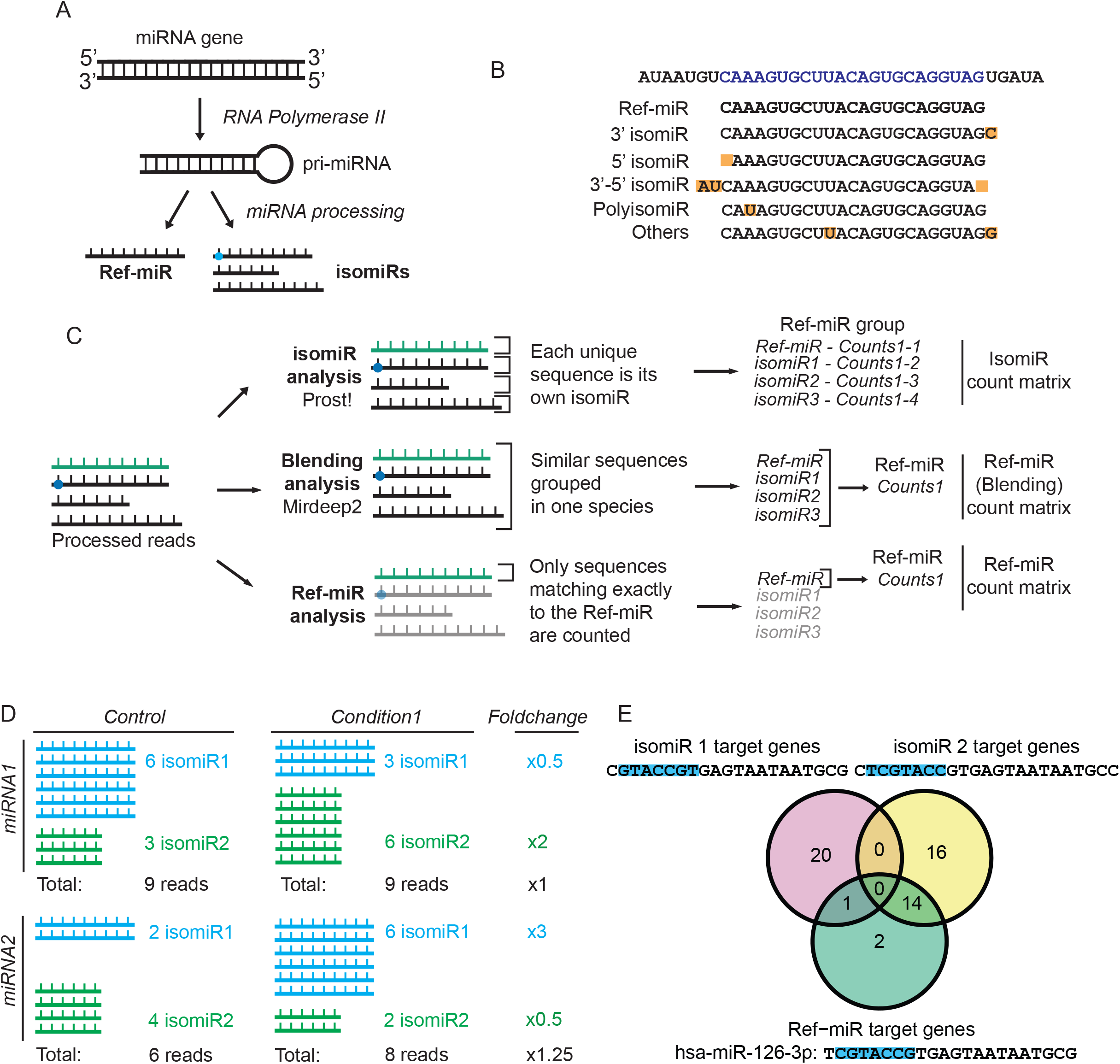
isomiR biogenesis, diversity, and potential impact on miRNA analysis. **A**. isomiR and miRNA biogenesis. A miRNA gene is transcribed into a primary miRNA (pri-miRNA) by the RNA Polymerase II. Various proteins, such as Drosha and Dicer, process the pri-miRNA further, cutting it into a mature miRNA sequence. In this process, alternative cleavage and additional modifications produce different isoforms (isomiRs) which differ from the reference sequence (Ref-miR). **B**. isomiRs can be categorised based on their alignment to the Ref-miR: *3’ isomiRs* and *5’ isomiRs* for 3’ and 5’ end modifications respectively. *3’-5’ isomiRs* for modifications in both ends. *PolyisomiRs* for single nucleotide modifications within the sequence. *Others* for all sequences that do not fit any of the previous categories (e.g. combination of these modifications). **C**. Three distinct methods of miRNA computational processing: isomiR analysis (reads are counted independently for each isomiR sequence), Blending analysis (summing up isomiRs from the same Ref-miR group) and Ref-miR analysis (only the sequences matching exactly to the Ref-miR are counted). These yields three distinct count matrices. **D**. When isomiRs are specifically regulated in a case-control experiment, Blending (in black) and isomiR (in blue and green) analysis can yield very different results. Adding up isomiRs in Blending analysis can potentially hide expression differences in isomiRs. **E**. Comparison of predicted targets for three miRNA species belonging to the miR-126-3p Ref-miR group (two isomiRs and the Ref-miR). Targets were predicted by inputing the seed sequence (in blue) of each miRNA in Targetscan’s seedmatch tool.

The discrepancy between evident isomiR importance and low amount of existing studies is the result of two main reasons. First, lack of a golden standard makes analysis and interpretation of the results challenging. Secondly, isomiRs are still considered as an optional addition to the standard miRNA analysis, as no study directly questions the validity of its results or clearly estimates the impact of isomiR exclusion. In a recent study (22), only half of the 133 differentially expressed isomiRs had also their reference miRNAs differentially expressed in standard miRNA analysis, while another study (23) shows isomiRs originating from the same reference sequence to be differentially expressed in opposite directions. This raises questions about the impact that isomiRs have on differential expression, the accuracy of the results, and the biological significance of the downstream studies.

Here, we assess the effects of isomiR inclusion on miRNA-seq results by using two Human Umbilical Vein Endothelial Cell (HUVEC) datasets. We determine the isomiR weight in terms of read quantity, added sequence diversity, and the evidence of sequence-specific expression changes. We then perform differential expression analysis using 1) isomiR miRNA analysis, 2) Blending (standard) analysis and 3) Ref-miR (reference miRNAs, no mismatch) analysis (**Fig 1C**) to explore and compare the results and investigate the potential biases and challenges introduced by the widely used approach that excludes isomiRs, and thus the consequences of ignoring isomiR existence in differential expression analysis (**Fig 1D**). Finally, we investigate condition-specific isomiR distributions, independently from global miRNA gene expression changes for higher resolution dissection of complex signals. The potential existence of isomiR specific differences and profiles could be of great importance as isomiRs have been shown to have different targets (24) (**Fig 1E**).

### Summary of the terminology

#### Ref-miR

The reference sequence of a given miRNA from the miRNA reference sequence database, miRbase. Synonym: canonical miRNA, miRNA reference sequence

#### isomiR

isoform of miRNA. isomiRs are defined relative to Ref-miR as they are classified based on the reference sequence alignment (**Fig 1B**). Ref-miR and Blending analysis both result in Ref-miR count matrices (**Fig 1C**), but ignore isomiRs, although Ref-miRs can be considered as an isomiR class (the canonical isoform of a miRNA).

#### Ref-miR group

A group of isomiR sequences that align to the same Ref-miR, and originate from the same miRNA arm. The Ref-miR group contains the Ref-miR. Ref-miR groups can also be called miRNa arm or miRNA species, but it is ambiguous as most miRNA studies do not account for isomiRs and use the term miRNA and miRNA arm as a synonym of the Ref-miR.

#### miRNA gene

The gene from which all isomiRs of two Ref-miR groups (3p and 5p arms) originate.

#### isomiR analysis

miRNA sequencing analysis method that counts isomiR reads independently, and annotates Ref-miR groups through alignment.

#### Blending analysis

Widely used analysis method for miRNA sequencing which adds up counts from sequences that align to the same Ref-miR, resulting in counts at the Ref-miR level only, blending potential isomiR signal. The analysis is based on the idea that biological diversity of miRNAs derives only from the Ref-miRs, and that isomiRs of the same Ref-miR group have the same signal and function. Also known as the canonical, classical, or standard analysis.

#### Ref-miR analysis

Ref-miR analysis only counts reads that are identical to the Ref-miR, discarding all isomiRs.

## MATERIAL AND METHODS

### Cell culture

For both the Senescence and compartment datasets, Human Umbilical Vein Endothelial Cells (HUVECs) were extracted with collagenase (0.3 mg/ml) digestion from umbilical cords obtained from the maternity ward of the Kuopio University Hospital. The collection was approved by the Research Ethics Committee of the Hospital District of Northern Savo, Kuopio, Finland. Written informed consent was obtained from the donors. The cells were cultivated in Endothelial Cell Basal Medium (Lonza) with recommended supplements (EGM SingleQuot Kit Supplements & Growth Factors, Lonza). Cells of seven donors were used in the study in separate, unpooled batches. All results were repeated on at least three donor batches.

### Sample preparation and miRNA sequencing

For the compartment dataset, HUVECs (passage 6) were cultured in T75 flasks in humidified CO_2_-incubator (0h control cells) or in a hypoxia chamber (Baker Ruskinn) with 1% O_2_, 5% CO_2_ for 7h or 24h. Cells were washed with PBS and collected by scraping to PBS+0.5% BSA. Cells were pelleted by centrifugation at 700g, +4 °C for 5 min and washed by PBS+0.5% BSA. Cell pellets were lysed with hypotonic lysis buffer and nuclear and cytoplasmic fractions were isolated according to the protocol by Gagnon et al. (25). Total nuclear and cytoplasmic RNA were extracted using TRIzol Reagent (Thermo Fisher Scientific) and RNA was dissolved into molecular biology grade water. Total RNA was treated with DNase I (cat. EN0521, Thermo Fisher Scientific) according to manufacturer’s instructions. RNA Clean & Concentrator-5 kit (cat. R1013, Zymo Research) was used to separate both nuclear and cytoplasmic total RNA into long and small RNA containing fractions according to the manufacturer’s protocol. RNA quality was assessed using Standard Sensitivity RNA analysis kit (cat. DNF-471-0500, Agilent Technologies) using Fragment Analyzer. Nuclear and cytoplasmic fraction separation was confirmed by qPCR for tRNA (htRNA-Lys-TTT-3-4). cDNA was synthesized using RevertAid Reverse Transcriptase (Thermo Scientific) and gene-specific primer (reverse primer) and quantified using Maxima SYBR Green/ROX qPCR Master Mix (2×) (Thermo Fisher Scientific). Thermal cycling was performed using a LightCycler480 (Roche) with the following program: 10 min at 95 °C, followed by 50 cycles of 15 s at 95 °C and 60 s at 60 °C. Primers used were (sequences are 5’ to 3’) Forward: GCCCGGATAGCTCAGTCG and Reverse: CGCCCGAACAGGGACTTG. The libraries were prepared using the NEBNext^®^ Multiplex Small RNA Library Prep Set for Illumina according to the instructions. Library sequencing was performed on the NextSeq 500 platform.

The sample preparation for the senescence dataset is described in (26). Briefly, HUVECs were extracted, with sample S0 corresponding to freshly isolated cells after collection, and S1, S2, S3 to first, second and third cell passage respectively. S0 samples represent the tissue-derived endothelial cells, which have grown in the presence of flow, and S1 to S3 samples are adjusted to static cell culture conditions. Of note, in standard HUVEC extraction, all endothelial cells originating from a given umbilical cord are plated and grown to confluence. In this experiment, only 29–36% of the harvested endothelial cells were plated for subsequent cell passages. Therefore, more population doublings were required from one passage to another compared to standard culturing, resulting in aged cells at S3 and cellular senescence by S6. The libraries were prepared using NEBNEXT library generation kit (New England Biolabs) according to the manufacturer’s instructions (the adapter sequence is AGATCGGAAGAGCACACGTCTGAACTCCAGTCAC). Samples were sequenced with the Illumina NextSeq 500.

### Preprocessing

The sequencing results were processed using bcl2fastq, provided by Illumina, and the resulting fastq files were further processed using fastxtoolkit. The command fastxclipper was used for adapter removal. The reads were then filtered by quality using fastq_quality_filter with parameters -q 30 and –p 90. The fastqc command was used to assess the quality of the sequencing.

### isomiR identification

isomiR sequences were discovered and characterized using *Prost!* Version 0.7.3 (27) according to the documentation of the program. The software was configured to only include sequences that had a total number of 25 reads across the two datasets, as well as a minimum length of 17 nucleotides and maximum of 25. Reads were aligned to the human genome GRCh 38 and the list of mature miRNA sequences and miRNA hairpins were obtained from miRBase (28). Target prediction of 3 species (2 isomiRs and their Ref-miR, hsa-miR-126-3p), was performed using TargetScan (29, 30). We imputed each species’s seed sequence in Targetscan Human Custom, v.5.2 (http://www.targetscan.org/vert_50/seedmatch.html).

### Blending analysis

As a conventional miRNA-seq processing pipeline, *miRDeep2* (31) was used to process the fasta files. From this pipeline, the command mapper.pl -c -j -m -s was run to collapse the reads and then quantifier.pl -p ../human_hairpin.fa -m ../hsa_mirna.fa -r -t hsa -d -j -y to align the sequences to the miRNA miRbase sequences (Ref-miRs) and their precursor. The resulting count files were then merged together. As miRDeep aligns not only to mature sequences, but also to precursors, some reads were counted twice and several rows of the same mature sequence existed in the counts table. This was corrected by removing duplicate rows.

### Ref-miR analysis

A Ref-miR specific count matrix was constructed by extracting the rows that correspond to sequences matching exactly to the Ref-miR from the *Prost!* isomiR count matrix. These counts thus originate from reads that are identical to the miRBase reference sequences.

### isomiR filtering and classification

From the *Prost!* output, sequence count matrix reporting the counts of each sequence and its alignment was extracted. Sequences that aligned to known transcripts (e.g. mRNA, tRNA, rRNA) but not to miRNAs were discarded. The rest were considered as “candidate isomiRs”. For all analysis, only sequences that aligned to the Ref-miRs, thus considered as isomiRs by *Prost!* were kept. isomiRs that *Prost!* aligned to multiple miRBase sequences were filtered out.

To categorize isomiRs, each isomiR was aligned to its corresponding Ref-miRs using Smith-Waterman local alignment method (32) (gap opening penalty -3, gap extension penalty -1), and then labeled accordingly: *Ref-miR* when the sequence was identical to the miRBase reference, *3’ or 5’ isomiRs* when it had an addition or deletion of nucleotide at the 3’ or 5’ end, respectively, but no other modifications, *polymorphic isomiR* when the isomiR retained the same length but one or more nucleotide changed, *3’-5’ isomiR* when it had an addition or deletion at both ends, and finally *other isomiRs* for other modifications or combinations.

In order to correct the counts for both sequencing depth bias and potential bias introduced by tRNA or rRNA levels, normalization was performed based on the total count of sequences that were considered as isomiRs by *Prost!*.

Sequences that had an average normalized count that was lower than a selected threshold (Reads Per Million, RPM) were discarded. For most downstream analyses, including differential expression, a threshold of 20 average RPM per sample was used. The expression level filtering was done independently for each dataset and applied to both blending and isomiR analysis.

### Differential miRNA expression analysis

For differential expression analysis, isomiRs were filtered independently for each dataset with a threshold of 20 RPM per sample (average). The miRNAs from the Blending analysis counts matrix were filtered similarly. For each condition, DESeq2 (33) was run independently on the non-normalized (raw) counts of all isomiRs (including miRBase sequences), the Blending analysis miRNA counts, and the Ref-miR counts matrix, independently. P-values were adjusted for FDR through DESeq2.

### Proportion and differential distribution analysis

Chi-Square test was performed using the chisq.test R function. isomiR proportion differences were explored by first calculating the proportion of each isomiR (dividing the number of counts for that isomiR to the total number of counts for the Ref-miR group). This was done independently for each sample. The aov R function was used to perform a one-way anova test for each isomiR and dataset to test if the proportion variance, within each isomiR independently, was linked to the sample group (the 4 senescence stage for senescence dataset, the 6 compartment-hypoxia combinations for the compartment dataset). Benjamini-Hochberg FDR correction was applied for the p-values, across all isomiRs.

## DATA AVAILABILITY

The compartment dataset has been deposited in NCBI’s Gene Expression Omnibus (34) and are accessible through [accession number] (link). The senescence dataset can be found at GSE94410 (https://www.ncbi.nlm.nih.gov/geo/query/acc.cgi?acc=GSE94410).

## RESULTS

### isomiRs are abundant, diverse and display sequence-specific patterns in their distribution

To determine the importance of isomiR inclusion in miRNA analysis, we first determined the global isomiR distribution characteristics across datasets, i.e. their abundance, added species diversity, and if there is evidence of non-random sequence-specific patterns.

To investigate the impact of isomiR presence in miRNA analysis, we compared three different methods to count reads (**Fig. 1C**): (1) *Prost!* for *isomiR analysis*, that groups isomiRs by their Ref-miR and counts all isomiR reads independently; (2) *miRDeep2* for *Blending analysis*, i.e. the most commonly used method in miRNA-seq analysis that aligns all similar reads (including isomiRs) and counts them as Ref-miR; and (3) *Ref-miR analysis* that only counts reads aligning to the Ref-miR without any mismatches or length variants.

We utilized miRNA-seq from two separate HUVEC datasets, namely Senescence and Compartment (**Fig 2A)**. The Senescence dataset was first presented in (26) and shows the effect of blood-flow (S0, n=3), and increasing senescence stages (S1, S2, S3, n=4 for each) on miRNA expression. The second is a novel dataset through which we explore miRNA expression under hypoxia (control, 7h and 24h of hypoxia, n=3), separately in the cytoplasmic and nuclear fraction (compartment-specific context).

**Figure 2:**
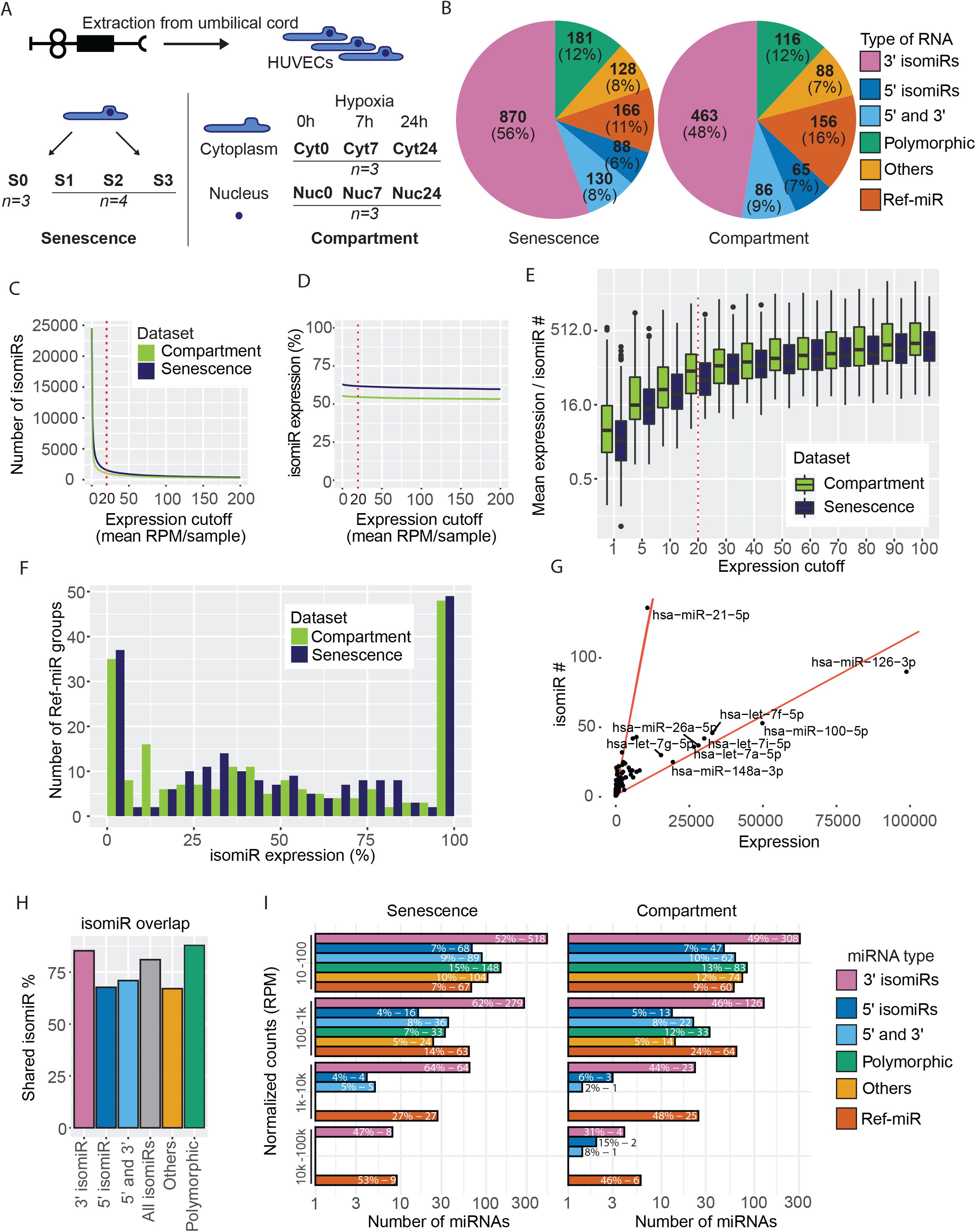
isomiR diversity in condition-specific HUVECs datasets. **A**. Description of the two datasets. **B**. Count distribution of isomiR categories in the datasets. Segments are colored by isomiR type. **C**. The number of detected isomiRs guides the choice of expression cutoff. The dotted red line marks the chosen cutoff (20 mean RPM). **D**. Percentage of miRNA expression stays stable with the expression cutoff. **E**. Distribution of the ratio between isomiR mean expression and the number of species, among different expression cutoffs. The ratio is calculated independently for each ref-miR group **F**. Distribution of reads (% originating from isomiRs, excluding Ref-miRs), amongst Ref-miR groups. **G**. Correlation between the number of isomiR species and average isomiR expression, for each ref-miR group, in the Senescence dataset. Two visual trends are marked in red lines. **H**. Percentage of shared isomiRs between the two datasets, using the compartment dataset as reference. **I**. Expression distribution of all miRNA species, for both datasets. Percentages are calculated within each expression group.

The results demonstrate that isomiRs account for the majority of reads, with many highly expressed species and distinct expression patterns (**Fig 2** and **Fig S1-4**). Both of the datasets had similar distributions of isomiR types (**Fig 2B** and **Fig S1**), and isomiR inclusion increased the miRNA diversity by several factors between the Blending and isomiR analysis (from 246 to 989 species in the Compartment dataset, and 279 to 1599 in the Senescence dataset).

A major challenge in isomiR analysis is that a vast majority of species are expressed at low levels and their biological relevance is uncertain. Previous studies have used various methods to select the isomiRs of interest (13, 35, 36). Most commonly this has consisted of a combination of low threshold and stringent selection of miRNA characteristics, such as number of allowed modifications or isomiR type. As isomiRs with the same seed sequence may also have specific biological effects (37–39), and very lowly expressed sequences are less likely to have strong biological effects, we filtered the isomiRs based on their expression levels only, using 20 RPM (mean per sample) as a cutoff **(Fig 2C-E)**. In both datasets, 50-60 % of the miRNA reads were isomiRs (the rest being Ref-miR reads), regardless of the chosen cutoff **(Fig 2D)**. Although the ratio of average expression per number of isomiRs increased with the cutoff, it remained relatively stable after 20 RPM **(Fig 2E)**.

For most Ref-miRs, a significant fraction of the total expression was exclusively originating from isomiRs (**Fig 2F)**. Only a small minority of Ref-miRs (less than 40) had low isomiR expression, and for some miRNAs, the Ref-miR itself was expressed below the threshold. In the senescence dataset, only 111 Ref-miR groups out of 279 in total had more than half of their expression coming from the Ref-miR sequence, the rest had most reads coming from isomiRs. Similarly, in the compartment dataset, only 116 Ref-miR groups out of 246, the majority of the expression originated from the Ref-miRs instead of isomiRs (**Fig S2**). Furthermore, the number of isomiRs for each Ref-miR was not solely dependent on the level of the miRNA expression (**Fig 2G and Fig S3**), suggesting sequence-specific isomiR generation, which created more isomiRs for specific sequences. This is the case for example for hsa-miR-21-5p, which had the highest number of isomiRs in the Senescence dataset although it ranked as 9th among the most expressed species (average expression per isomiR).

isomiRs that were detected above the threshold were mostly shared between the datasets (**Fig 2H**). The percentage of shared isomiRs remained above 75% regardless of the selected cutoff (**Fig S4**). Overall, the high similarity between the miRNA species in the two separate datasets suggests that isomiR generation is not a random event but a well-regulated, sequence-specific process.

A more detailed breakdown of expression level distribution **(Fig 2i)** among isomiR types showed that individually, some isomiRs were among the highest expressed species in both datasets (10K - 100K RPM: eight and seven isomiRs for the Senescence and Compartment datasets, respectively). We also observed heterogeneous expression levels between the isomiRs types, with no *Polymorphic* and *Other* isomiRs (isomiRs that do not fit the categories of **Fig 1B**, e.g. combination of the categories) seen above 1,000 RPM.

In some cases, the Ref-miR to which the isomiR aligns to is not expressed in the dataset above the detection threshold, though these cases are a minority (not more than 25% of Ref-miR groups). Interestingly, in the *Polymorphic* and *Others* isomiR category, the Ref-miR is expressed in all cases. These categories also have a high number of isomiRs relative to the number of Ref-miR groups (ratio of number of isomiRs / number of Ref-miR groups is over nine against three for 3’ isomiRs).

Interestingly, these polymorphic isomiRs, in both datasets, belong to the same Ref-miR group (10 out of 11, **Suppl. Table 1**). These 10 Ref-miR groups are hsa-miR-100-5p, hsa-miR-22-3p, hsa-miR-10b-5p, hsa-miR-21-5p, hsa-miR-381-3p, hsa-miR-126-3p, hsa-let-7i-5p, hsa-let-7f-5p, hsa-let-7g-5p, hsa-miR-26a-5p. All of these particularities suggest highly sequence-specific post-transcriptional modifications for isomiR generation, resulting in different distribution properties.

The multiple sequence specific expression patterns that we highlight here are concordant with tight regulation of the species and potential effect of cellular context (condition or compartment) in their expression. Coupled with their high abundance and diversity, this makes isomiR differential regulation and expression across conditions very likely.

### isomiR-level analysis show major changes in expression and discrepancies with Blending analysis

We next asked if isomiRs were differentially expressed between conditions, and if so, how does the differential expression impact the reliability of Blending analysis, which groups reads by Ref-miR. To answer the first part of the question, we used isomiR analysis to determine differential expression results at the isomiR level. The analysis identified a vast number of significantly differentially expressed isomiRs, with log fold changes varying from 10 to -5, and very high significance (**Fig 3A**). For most conditions, all isomiR types were represented among differentially expressed miRNAs **(Fig S5)** and some of the differentially expressed isomiRs were shared across comparisons. For example, 266 isomiRs of the S3 to S0 comparison were also found to be differentially expressed in the S1 to S0 comparison, including flow responsive miRNAs that are known to participate in the regulation of flow-dependent angiogenesis, such as isomiRs for hsa-miR-31-5p and hsa-miR-100-5p (26) (**Fig 3B**). Over 600 isomiRs were differentially enriched between the nuclear and cytoplasmic compartments, out of 974 detected species (**Fig S6B**). Both in the Nuclear to Cytosolic and S3 to S0 comparisons, 60% of isomiRs were found to be significantly differentially expressed, which is a comparable percentage to the Blending and Ref-miR analyses (**Fig S6A**), supporting a hypothesis that isomiR analysis is not noisier than Blending analysis. Furthermore, for many miRNAs, the coefficient of variation was lower for isomiR than for Blending analysis (**Fig S7**). Overall, the greater number of isomiRs compared to Ref-miRs resulted in up to 5 times more differentially expressed species in IsomiR analysis compared to Ref-miR analysis (**Fig S6B-C**).

**Figure 3:**
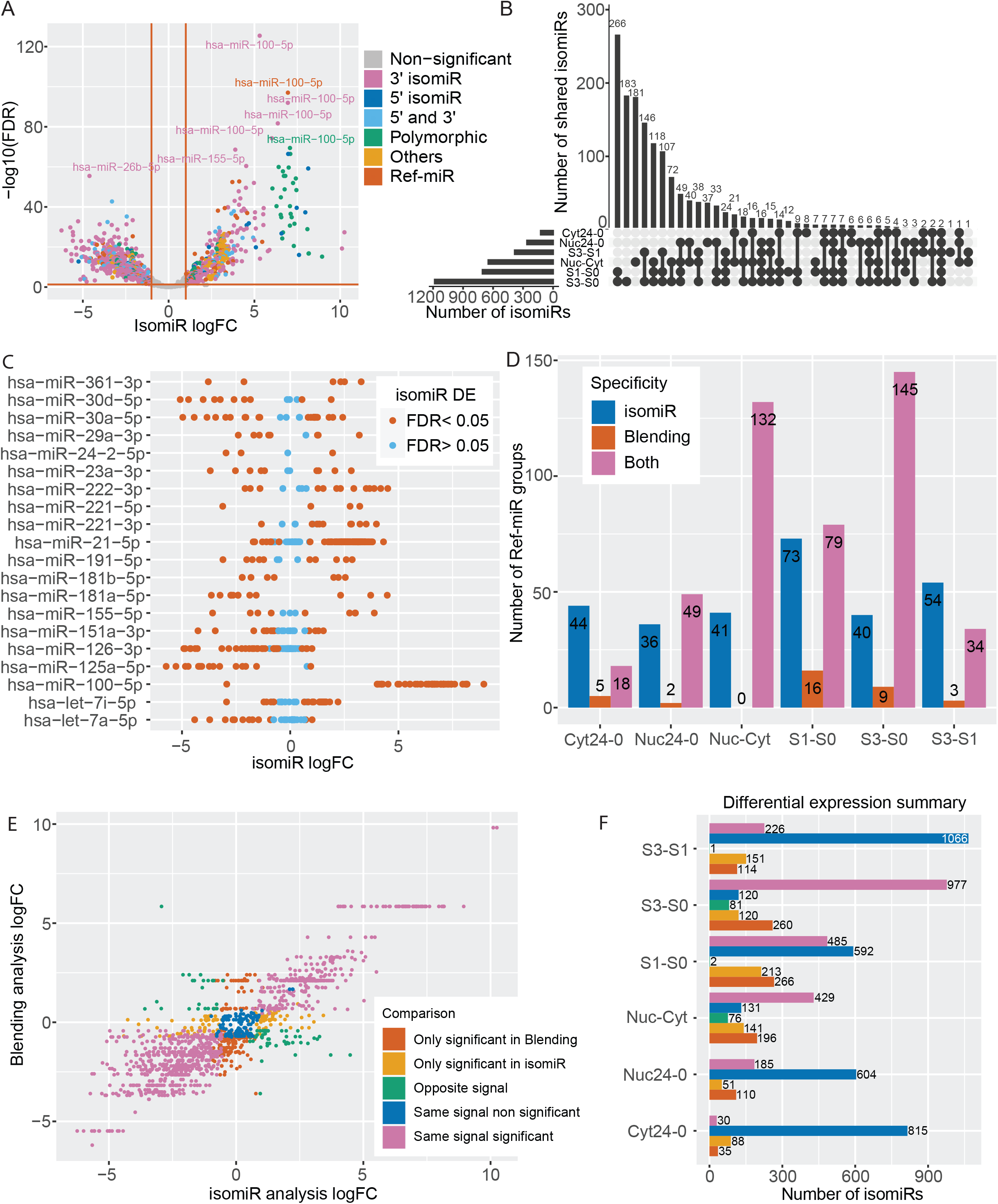
Comparison of differential expression results from isomiR and Blending analyses. **A**. isomiR differential expression (DE) results for S3-S0 comparison in the Senescence dataset. **B**. Upset plot showing intersection of significantly DE (FDR < 0.05) isomiRs across datasets and comparisons. **C**. isomiR logFC distribution in the S3-S0 comparison, for the top 20 Ref-miR groups with the highest variation in their isomiR logFC. Each dot is an isomiR from the line’s ref-miR group, colored by significance. **D**. Distribution of DE results per Ref-miR group, across comparisons and datasets. Ref-miR groups are arranged in bins: *isomiR* (Blue) is when one isomiR of the Ref-miR group is DE at the isomiR level only (one of the Ref-miR group’s isomiRs), and *Blending* (red) when the Ref-miR is DE in Blending analysis only. *Both* (pink) are cases where the Ref-miR group is DE in both Blending and isomiR analysis. The Nuc-Cyt comparison includes all nuclear and cytoplasm samples. **E**. LogFC comparison between isomiRs and their associated Ref-miR groups, for S3-S0 comparison, within the Senescence dataset. Each dot is an isomiR, colored by comparison group: *Only significant in Blending* means that the isomiR is not significantly DE but its Ref-miR group is DE in Blending analysis. *Only significant in isomiR* are for species for which Blending analysis yields no significant signal, but that are DE in isomiR analysis. When both the isomiR and its Ref-miR are DE but with opposite logFC, the situation is classified as *Opposite signal*. Finally, the last cases are non DE in both methods (*Same signal non significant*) or DE in both, with the same direction (*Same signal significant*). **F**. Results summary of DE in isomiR and Blending analyses across comparisons and datasets. The barplot shows the number of isomiRs in each comparison category (sames classes and colors as in E).

To address the second part of the posed question, we further compared the results of the isomiRs to Ref-miR results to estimate the potential effect of the Blending approach on result accuracy. For some Ref-miRs, the fold changes of their isomiRs varied greatly (**Fig 3C**), showing differentially expressed isomiRs with opposite fold changes, in addition to isomiRs with fold changes close to 0, all for the same Ref-miR. Similar results were observed for all compared conditions (**Fig S8**). This heterogeneity in the context-specific isomiRs puts in question the Blending analysis principle of binning all species from the same Ref-miR together, as they can display very different signals.

In both datasets, 18 to 36% of Ref-miRs were not differentially expressed in Blending analysis but had at least one isomiR that was differentially expressed (Fig 3D). This number exceeded the inverse case where differential expression was observed in Blending analysis but not in isomiR analysis by up to 28 percent of Ref-miRs. For example, in the S1 to S0 comparison (Senescence dataset), 79 Ref-miRs contained at least one differentially expressed isomiR and were also differentially expressed in Blending analysis themselves, while only 16 miRNAs were not significant in isomiR analysis while being differentially expressed in the Blending analysis, and 73 miRNAs that were not significant in the Blending analysis were found to be differentially expressed at the isomiR level, highlighting the information loss in Blending analysis.

Most Ref-miRs contained isomiRs with a wide range of fold changes, in addition to discordances between IsomiR and Blending analysis as shown for the S3 to S0 comparison, in Fig 3E and for the other conditions in Fig S9. In some cases, isomiRs were significantly differentially expressed, but in a different direction than their Ref-miRs, which were also significantly differentially expressed in the Blending analysis. In the S3 to S0 comparison, there were 81 such cases (Fig 3F). Consistently, a high number of miRNAs (213 in S1 to S0 comparison) were significantly differentially expressed at isomiR level but not in Blending analysis, meaning that this signal would be lost without isomiR level analysis. Beyond discrepancies, isomiR analysis also permits to pinpoint, within a differentially expressed Ref-miR group, which isomiRs are differentially expressed, and which are not, improving the resolution.

The broad and diverse differential expression signal shown here suggests major differential distribution in isomiRs between biological conditions, with different signals between isomiRs from the same Ref-miR group, contradicting the principles of Blending analysis. This is supported by the many discrepancies between IsomiR and Blending analyses, and the added information and resolution that IsomiR analysis provides.

### Ref-miR-specific analysis further highlights the limits of the Blending analysis

Ref-miR analysis, which allows no variation to the reference sequence, is very rarely used in miRNA studies. Nonetheless, we explore this third miRNA processing method and compare it with isomiR and Blending analysis to further study the impact of isomiRs on Blending analysis results. Notably, Blending analysis is based on the hypothesis that sequences similar to the reference sequence have the same function and expression distribution, and thus can be binned together under the reference sequence.

Our results showed that the Ref-miR differential expression signal further differed from the isomiR signal compared to the Blending analysis (**Fig 4A-B and Fig S10**). This suggests that a Ref-miR and its isomiRs can have very different responses to stimuli, and therefore questions the standard way of analysing miRNAs by grouping the isomiRs under reference sequences and adding all counts together (Blending analysis). The analysis revealed many Ref-miRs that were not differentially expressed but one of their isomiRs was (**Fig 4C**).

**Figure 4:**
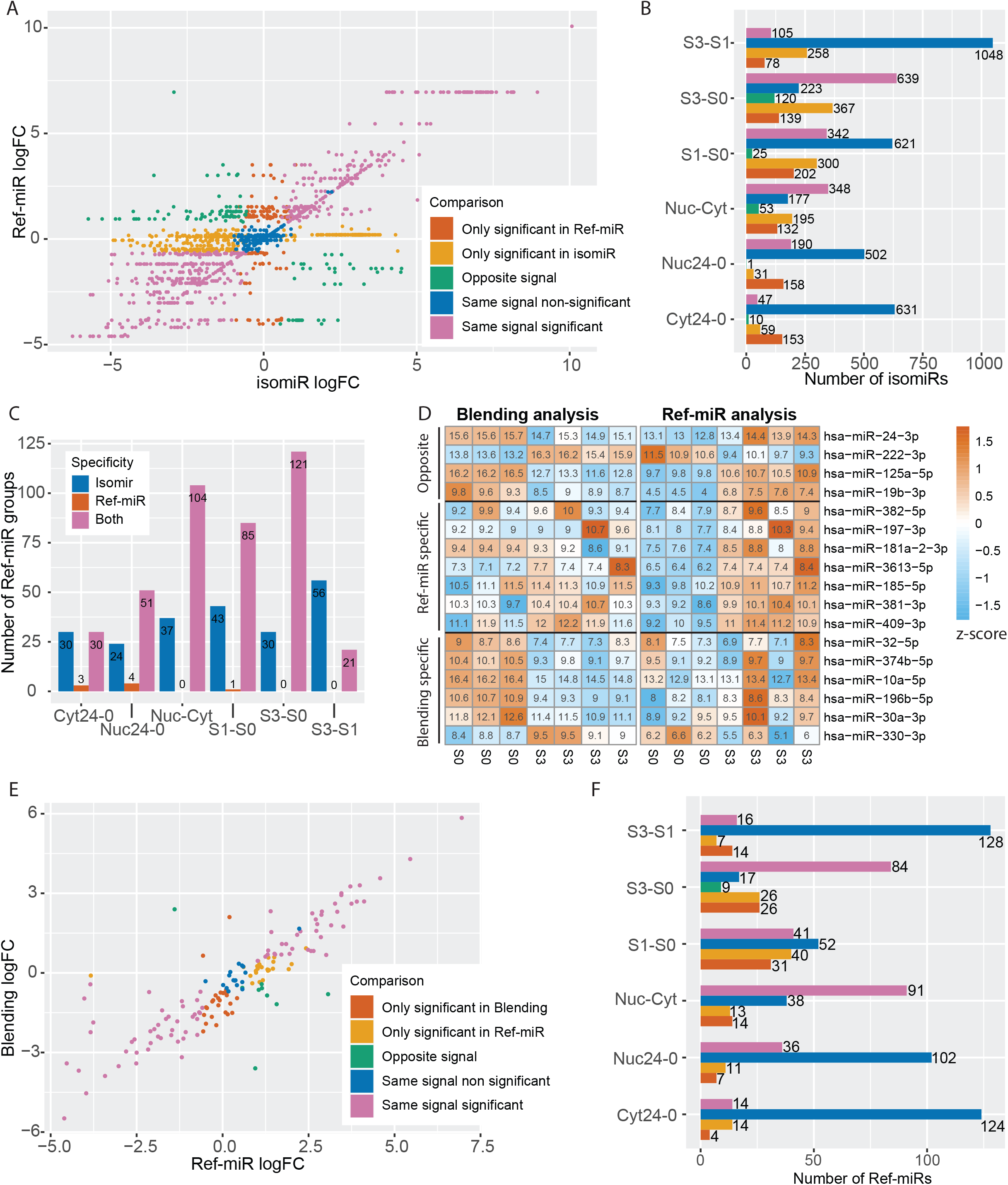
Ref-miR specific DE analysis. **A**. LogFC comparison between isomiR and Ref-miR analysis groups, for the S3-S0 comparison, in the Senescence dataset. Each dot is an isomiR, colored by comparison group (groups described in 3E, with Ref-miR analysis replacing Blending analysis). **B**. Results summary of DE in isomiR and Ref-miR analyses across comparisons and datasets. The barplot shows the number of isomiRs in each comparison category. **C**. Distribution of DE results per Ref-miR group, across comparisons and datasets. Ref-miR groups are arranged in bins: *isomiR* (Blue) is when one of the Ref-miR group’s isomiR is DE at the isomiR level only, and *Ref-miR* (red) when the Ref-miR is DE in Ref-miR analysis only. *Both* (pink) are cases where the Ref-miR group is DE in both Ref-miR and isomiR analysis. **D**. Heatmap of Deseq2 normalized expression in the Senescence dataset (S3-S0). miRNAs of interest are shown, comparing Blending and Ref-miR results for the same miRNAs. miRNAs are categorized as *Opposite signal, Ref-miR specific* (only DE in Ref-miR analysis) and *Blending specific* (Only DE in Blending analysis). **E**. LogFC comparison between Blending analysis and Ref-miR analysis, for S3-S0 comparison, within the Senescence dataset. **F**. Summary of DE in Ref-miR and Blending analyses across comparisons and datasets.

The Blending hypothesis implies high similarity between the Ref-miR and the Blending analysis results, nevertheless, we observed many discrepancies, similar to previous comparison between Blending and IsomiR analyses. In some cases, miRNAs had significant but opposite signals (**Fig 4D**). For example, hsa-miR-24-3p was downregulated in senescence while the reference sequence on its own was upregulated based on Blending analysis and Ref-miR analysis, respectively. This was the case for nine other isomiRs (**Fig S11**), and the signal was reversed for hsa-miR-222. For all comparisons, there were cases where the miRNA was significantly differentially expressed in Blending analysis but not in the Ref-miR analysis, and vice versa. In some of these cases, there was a clear trend toward the opposite signal, although non-significant (e.g. hsa-miR-374, hsa-miR-10a, hsa-miR-196b on **Fig 4D**). Even Ref-miRs that displayed the same direction in differential expression between the two methods could vary in their fold changes, as shown in **Fig 4E** and **Fig S12**, suggesting additional signal differences that may be biologically relevant.

These cases were not merely anecdotal examples, as in the S1 to S0 comparison, there were almost twice as many miRNAs that had diverging signals than miRNAs with significant signals in the same direction (**Fig 4F)**. Most of these cases arose from the situation where the signal captured by the Blending analysis originated from isomiRs instead of the Ref-miR miRNA, or where the sum of all isomiRs for the same Ref-miR drowned the Ref-miR-specific signal.

Taken together, as opposed to the Blending analysis hypothesis, we showed major discrepancies between Ref-miR and isomiR analysis results. These results highlight the problems in the Blending analysis and questions the biological relevance of the follow-up studies conducted based on the Blending analysis, when the signal in the analysis is interpreted to result from the expression changes of the reference sequence only.

### isomiRs allow for in-depth exploration of specific biological conditions

Having established the significance of isomiR-level analysis and its promise of higher resolution results, the question arose as to what extent such analysis could provide information on condition-specific miRNA regulation. To this end, we explored isomiR response to complex signals through the Compartment dataset, which represents a combination of nuclear and cytoplasmic miRNAs under normoxia and hypoxic conditions.

IsomiR-level analysis of the compartment-specific data revealed that the majority of species were shared across all comparisons, and thus were differentially expressed regardless of hypoxia levels (**Fig 5A**). In total, 107 miRNAs were differentially expressed between compartments in hypoxic conditions only, while 137 miRNAs were differentially expressed between compartments in normoxic conditions only. Similarly, there was a clear difference between the cytoplasmic and nuclear compartments for differential expression of hypoxia-responsive isomiRs (**Fig 5B**). 150 isomiRs were differentially expressed in the nucleus, both at 7 and 24 hours of hypoxia, but not in the cytoplasm, and 78 isomiRs were differentially expressed in 24h of hypoxia, in the cytoplasm only. Furthermore, there globally was more differentially expressed isomiRs in hypoxic conditions in the nucleus than in the cytoplasm, and almost no differentially expressed isomiRs at 7 hours of hypoxia in the cytoplasm. 55 isomiRs were differentially expressed with hypoxia in both the nucleus and cytoplasmic compartments, while some other isomiRs were differentially expressed only in hypoxia at specific stages (76 species at 24 h in the nucleus, and 60 at 7 h). Consistently in the senescence dataset, we observed 595 isomiRs being uniquely differentially expressed in freshly isolated samples compared to those adjusted to static cell culture conditions (**Fig 5C**). Moreover, beyond the compartment-specific differential expression, we observed that the isomiR type distribution was significantly different between the nucleus-enriched and cytosol-enriched species (p <1.80e-07 in X^2^ test, **Fig 5D**), suggesting differential regulation of isomiR sequences, as nuclear-enriched isomiRs may be implicated in non-canonical regulatory functions in the nucleus (40, 41).

**Figure 5.**
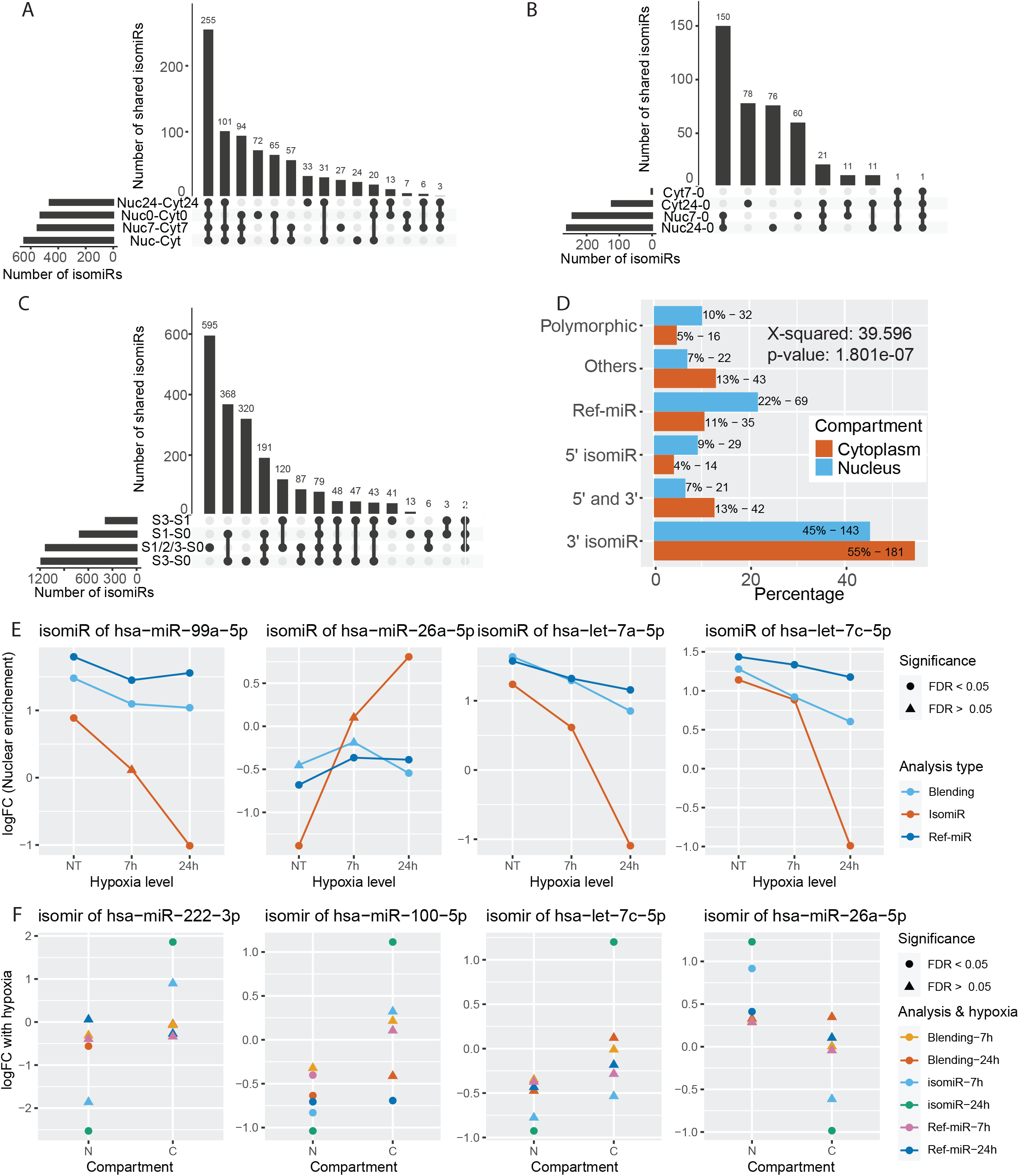
Response of isomiRs in complex biological conditions. **A**. Upset plot of compartment enriched isomiRs across hypoxia stages. **B**. Upset plot of hypoxia DE isomiRs across compartments and hypoxia stages. **C**. Upset plot of Senescence DE across conditions. **D**. Distribution of categories for isomiRs that are DE between compartments (Nuc-Cyt). Statistical dependance of isomiR type and compartment enrichment is assessed using a X-squared test (p-value = 1.801e-07). **E**. Effect of hypoxia on compartment enrichment for four isomiRs and their Ref-miRs. The colors correspond to the type of analysis (light blue for Blending analysis, red for isomiR analysis and dark blue for Ref-miR). Dots are used for significant DE and triangles for non significant results. **F**. Differential expression results for four isomiRs and their Ref-miRs (0h vs. 7h and 24h of hypoxia), within compartments, using Blending analysis (orange for 7h and red for 24h), Ref-miR analysis (purple for 7h and dark blue for 24h) and isomiR analysis (light blue for 7h and green for 24h). Dots are used for significant DE and triangles for non significant results.

Next, we set out to explore the impact of hypoxia on the compartment-specific isomiR abundance **(Fig 5E)**. A total of 11 isomiRs and 7 Ref-miRs were differentially enriched between nucleus and cytosol in response to hypoxia, with isomiR and Ref-miR analysis respectively, but not in Blending analysis (**Fig. S13**). For example, hsa-miR-99a-5p was significantly enriched in the nucleus compared to cytosol both in normoxia and under hypoxic conditions according to the standard and Ref-miR analysis, whereas one of its isomiRs was significantly enriched in the nucleus in normoxia, but switched to cytoplasm at 7 hours of hypoxia. Similarly, an isomiR of hsa-miR-222-3p was upregulated with hypoxia (24h vs. 0h) in the nucleus and depleted in the cytoplasm (24h vs. 0h of hypoxia), while its Ref-miR showed no significant changes either in Blending or Ref-miR Analysis (**Fig 5F**). Altogether, we identified 11 cases showing compartment-dependent responses to hypoxia (9 isomiRs and 2 Ref-miRs, **Fig 5F and Fig S14**), indicating existence of hypoxia-dependent isomiR transport between compartments, and compartment-dependent isomiR expression. Furthermore, many of these complex, statistically significant signals were not detected in the Blending analysis.

Taken together, the results indicate that isomiR-level analysis permits high-definition dissection of condition-specific signals, as these signals are highly affected by isomiR dynamics. The observed changes, seen at multiple levels, such as intersection of differential expression signal, differential distribution of isomiRs types and interactions between conditions, further enforces the importance of isomiR analysis.

### isomiRs are differentially enriched irrespective of global miRNA signals

In this study, we showed that isomiRs display novel condition- and compartment-specific expression patterns through differential expression analysis. However, isomiR expression levels depend both on their primary miRNA transcription as well as the regulation of isomiR biogenesis (**Fig 1A**). Therefore, we next set out to decipher whether the differential signal in our datasets arises from: 1) primary transcription, which would affect all isomiRs and their Ref-miR signal, leading to isomiRs exhibiting differential expression without any changes in the distribution within their Ref-miR groups (**Fig 6A - case 1**), or from 2) isomiR biogenesis, which would be expected to cause changes in distribution, despite the same global signal, suggesting a condition specific regulation of isomiRs independently of the Ref-miR group signal and miRNA transcription (**Fig 6A - case 2**), by using isomiR proportions as an independent metric of isomiR levels.

**Figure 6:**
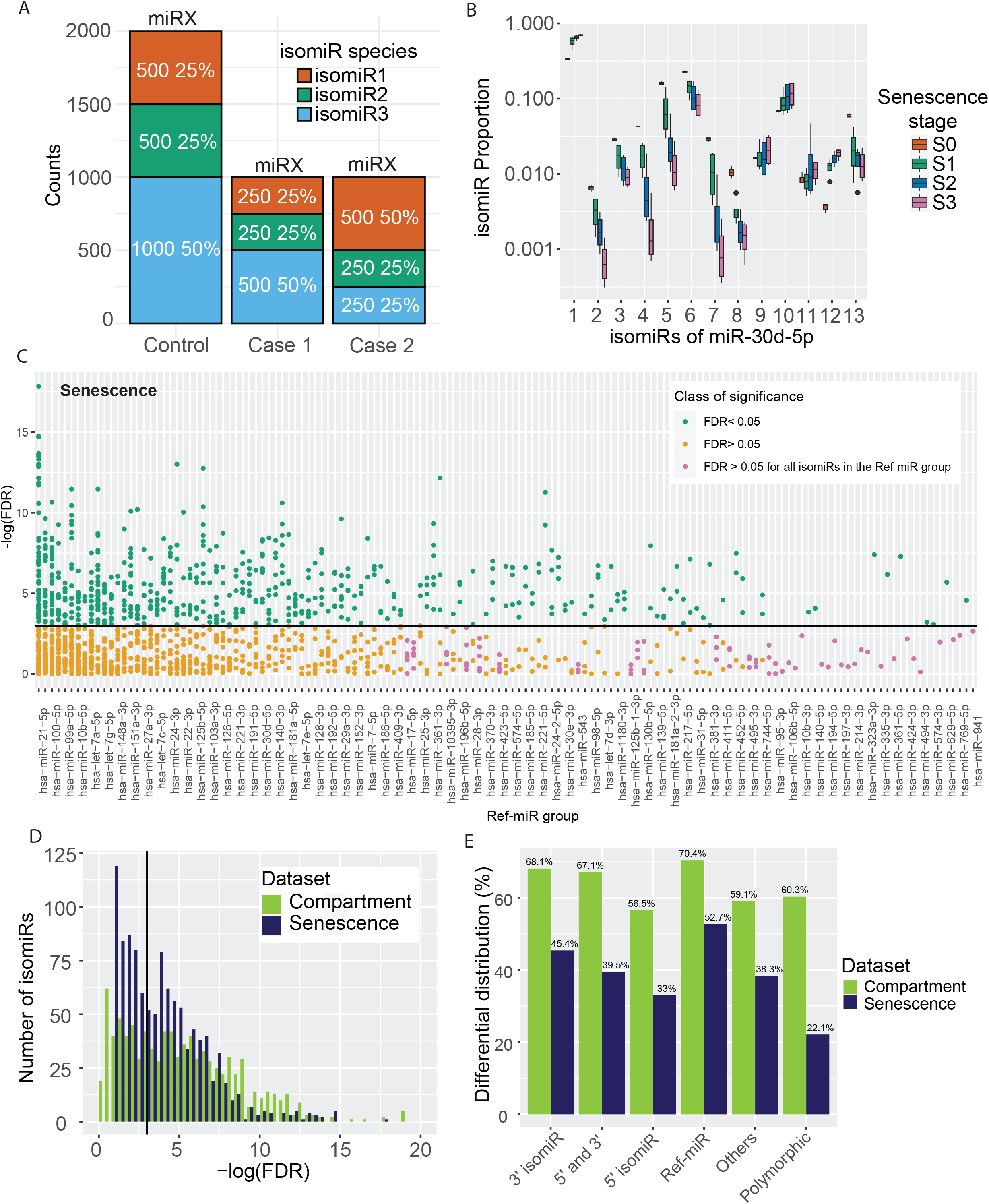
Differential distribution (DD) of isomiRs using proportion based analysis. **A**. miRNA gene expression and isomiR regulation impacts isomiR differential expression analysis. In case 1, all isomiRs are DE, but their proportions are the same. In case 2, there is an overall decrease of isomiRs expression levels, in addition with a change of proportion, suggesting isomiR specific response to biological conditions, independently of the global miRNA transcription signal. **B**. Distribution of isomiR proportions for hsa-miR-30d-5p, across senescence stages. **C**. Breakdown of isomiR differential distribution analysis. For each Ref-miR group, we show the statistical significance level of its isomiRs differential distribution results. Species that are significantly DD are shown in green, the ones that are not DD in orange and species for which no isomiR in the Ref-miR group is significantly DD are shown in purple. **D**. Distribution of proportion-based Anova adjusted p-values among isomiRs, in both datasets. Significant result on the Anova test indicates that the isomiR species is differentially distributed. **E**. Percentage of DD isomiRs between categories, for both datasets.

We started by exploring isomiR proportions between conditions using hsa-miR-30d-5p isomiRs as an example. Our analysis revealed low proportion variance within the same senescence stage but high variance between the different stages, illustrating clear proportion differences between conditions and thus specific isomiR distribution changes (**Fig 6B)**. In order to generalize the results to all isomiRs, we used an Anova test with FDR correction on the isomiR proportions. In the Senescence dataset, 619 out of 1599 detected isomiRs were differentially distributed (i.e., their proportions changed between conditions, **Fig 6C, 6D and S15**). Interestingly, most Ref-miR groups contained at least one isomiR that was differentially distributed, and the ones that did not, usually only had a few isomiRs. Differentially distributed isomiRs were thus dispersed among Ref-miR groups and not concentrated in a few of them (**Fig S16A**). Similar distributions could be observed in the compartment dataset, where 596 isomiRs out of 989 were differentially distributed (**Fig 6D, S15A, and S16A**). The strength of the observed differential distribution signal is supported by the wide range of percentages of differentially distributed isomiRs that is present within each Ref-miR group, going up to 100% for 50 species in the Compartment dataset (**Fig S16B**). Only a minority of Ref-miR groups, (20-40 in total for 240 groups) have no differentially distributed isomiRs.

Of note, isomiR categories had different proportions of differentially distributed species (**Fig 6E**). The proportions ranged from 20 to 52 % in the senescence dataset, and 56 to 70 % in the compartment dataset, suggesting a functional specificity for the different types of modifications that generate isomiRs. In line with these results, differences in variance for condition-dependent isomiR proportions were also observed between categories (**Fig S17**).

Altogether, our results suggest that isomiRs display wide proportion differences, for an important fraction of isomiRs, Ref-miR groups, and isomiR types. These results confirm the strength of the observed differential expression signal, its wide presence across miRNAs, and independence from global miRNA transcription signal, supporting the condition-dependent isomiR regulation that was inferred from differential expression analysis.

## DISCUSSION

In this study, we show evidence to support the systematic, in-depth, isomiR analysis as a complement or replacement for Blending analysis. We highlight the problems involved in Blending analysis that can result in largely incomplete and even incorrect results. We confirm that isomiRs are present in high numbers with sequence-specific distributions and high diversity. They are systematically differentially expressed in several conditions, and their differential expression signal diverges significantly from Blending analysis, in many cases increasing sensitivity. Furthermore, isomiR analysis permits the exploration of complex interactions between different biological contexts, providing a more complete picture of miRNA profiles across biological states. Moreover, we show that isomiRs are broadly differentially distributed, independently from the global miRNA signal, providing evidence of isomiR-specific regulation in different conditions.

Nevertheless, there are also caveats and limits to consider when performing isomiR analysis. The studied sequences differ from each other by only one or two nucleotides, which supports counting isomiRs without alignment-based bins, but through exact sequence counts, making them vulnerable to, for example, sequencing errors. The high similarity in sequence diversity in both of our datasets, in addition to high cutoff thresholds, enabled us to limit the effect of potential sequencing errors on our results. However, as both datasets were generated with the same library preparation methods, some of the diversity could be the result of library-specific bias, which have been shown to impact isomiR analysis (42). While more exploration is warranted, for example using paired-end sequencing (43), this bias cannot explain differential expression and distribution results, as it is not condition-specific in nature.

Computational analysis of isomiRs remains difficult, as no golden standard exists for their identification, stringency for isomiR alignment and selection is not clear, and expression cutoff remains arbitrary. Some methods only allow for specific modifications and isomiR types, with a limited number of potential nucleotide changes, such as Seqbuster (35, 44). Some methods use alignment on the whole genome (27), while others are Ref-miR alignment based (45). As isomiR signal results from miRNA gene expression and isomiR biogenesis, new modeling based methods that take both into account would be beneficial, thus combining the differential expression and differential distribution methods presented here. Previous studies that evaluate isomiR expression response only used differential expression analysis, not correcting for global miRNA expression changes (11, 22, 46). Our study provides a pipeline of comprehensive miRNA analysis, for differential expression in Blending, Ref-miR, and isomiR analysis, in addition to differential distribution, ensuring the capture of the most signal from miRNA sequencing to date.

To avoid skewing the results, we removed ambiguous sequences that could be aligned to more than one Ref-miR. The existence of such sequences is problematic for the Blending analysis, where these reads are usually split between Ref-miRs and the binned reads for each Ref-miR are seen as the overall signal for the Ref-miR in question. To clarify their biological significance, these ambiguous sequences should be further explored. Some miRNA analysis methods try to solve ambiguous sequence alignment using quality scores (47, 48), but it doesn’t solve the case of totally ambiguous isomiRs for which there would be no possible basis to assign them to one reference sequence or another.

As canonical function of miRNA species involves the target hybridisation with the seed sequence, 5’ modifications can easily modify the target and the function of the isomiR compared to its Ref-miR, as is supported by many studies (49–52). Nevertheless, other isomiR types, especially the highly diverse group of 3’ isomiR, can also be of functional importance, as the 3’ end composition affects Argonaute 2 affinity for canonical miRNA function (53), and non-canonical functions are more and more described for miRNA species (40, 41) including in the nucleus, where our analysis shows specific enrichment of isomiR species and types. Functional importance of 3’ modifications is further supported by previous studies (37–39). In order to confirm and explore the scale of importance of the presented results, more isomiR functional studies are called for, to have a clearer view of their functional importance. Especially, isomiR-specific target prediction softwares or databases would be of high interest, as most target prediction algorithms are suited for mature Ref-miRs only. Such tools could use both prediction algorithms and target discovery experiments to infer isomiR function, as studies support non-seed target recognition (54), and non-canonical target-gene regulation (55).

While the Senescence dataset is taken from a previous study (26), with only superficial isomiR analysis, the Compartment dataset is novel, thus providing a resource of compartment-specific miRNA expression in hypoxic conditions. This dataset highlights that isomiRs and miRNA species are expressed at different levels in the nucleic or cytosolic fractions, and that hypoxia significantly modulates their expression level. In addition, we show differences in isomiR hypoxia response between compartments, and changes in the compartmental enrichment of some isomiRs depending on the hypoxia timepoints. This suggests hypoxia related functions for these species, both in the nucleus and cytoplasm, with specific regulatory mechanisms.

In conclusion, our results provide a novel framework for miRNA analysis that takes into account isomiR expression dynamics. We show that ignoring isomiRs could result in missing most of the species diversity, omitting a great amount of differential expression and distribution signal, and misrepresent the reality by summing up reads from various sequences that display different or even opposing expression signals. Instead of the widely used Blending analysis, we advocate for general inclusion of isomiRs in all miRNA sequencing analysis, and isomiR-level differential expression and distribution analysis, to fully capture the information given by miRNA sequencing.

## ACKNOWLEDGEMENTS

We are grateful to Tuula Salonen for excellent technical assistance in library preparation. We acknowledge the University of Eastern Finland (UEF) Genome Center and Biocenter Finland for infrastructure support. We acknowledge Carles A. Boix and Pierre R. Moreau for their discussion and insight in this study.

---

**Supplementary Figure 1:**
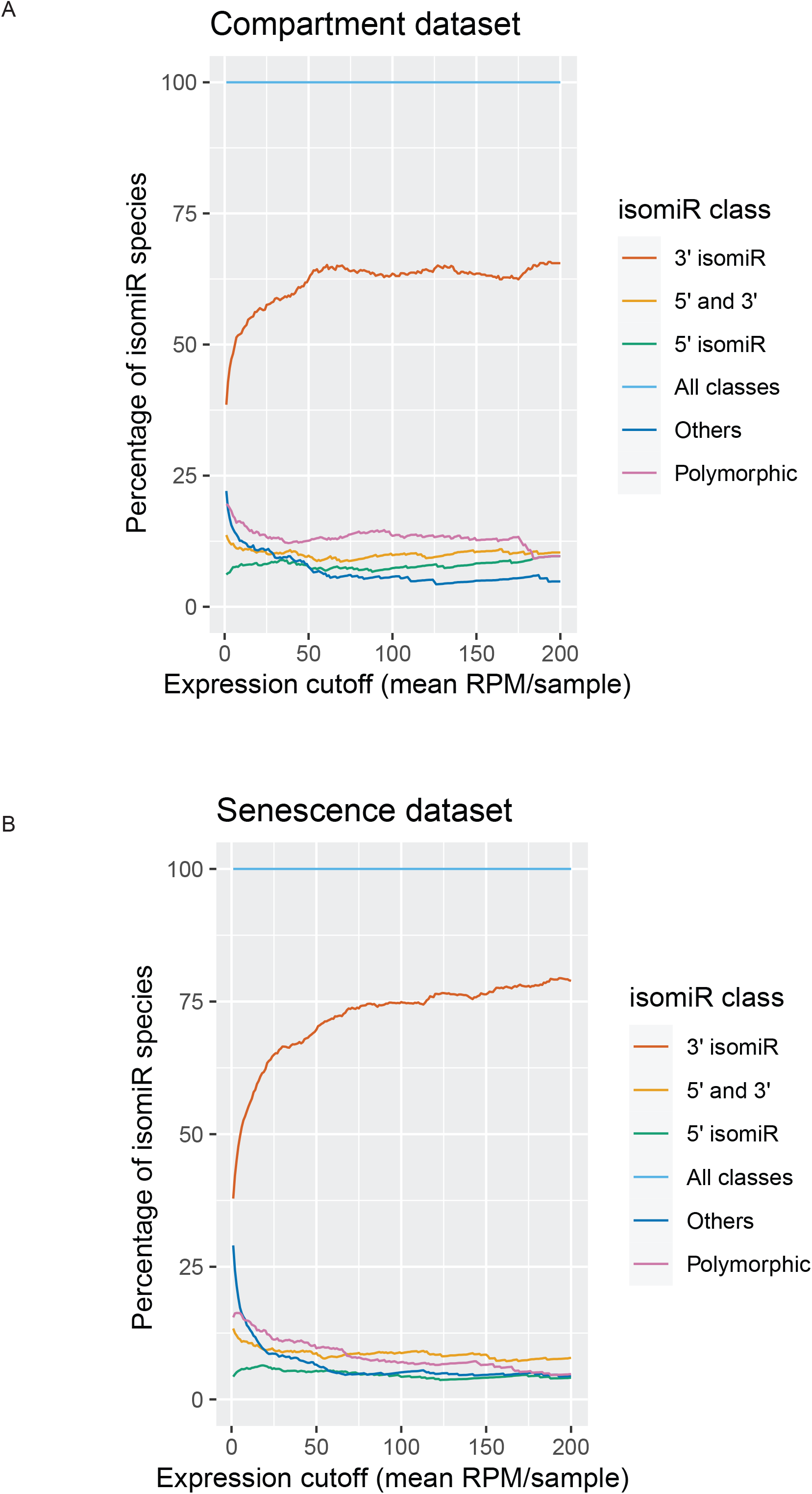
Distribution of isomiR species types depending on the expression filtering cutoff, in. **A**. Compartment dataset and **B**. Senescence dataset.

**Supplementary Figure 2:**
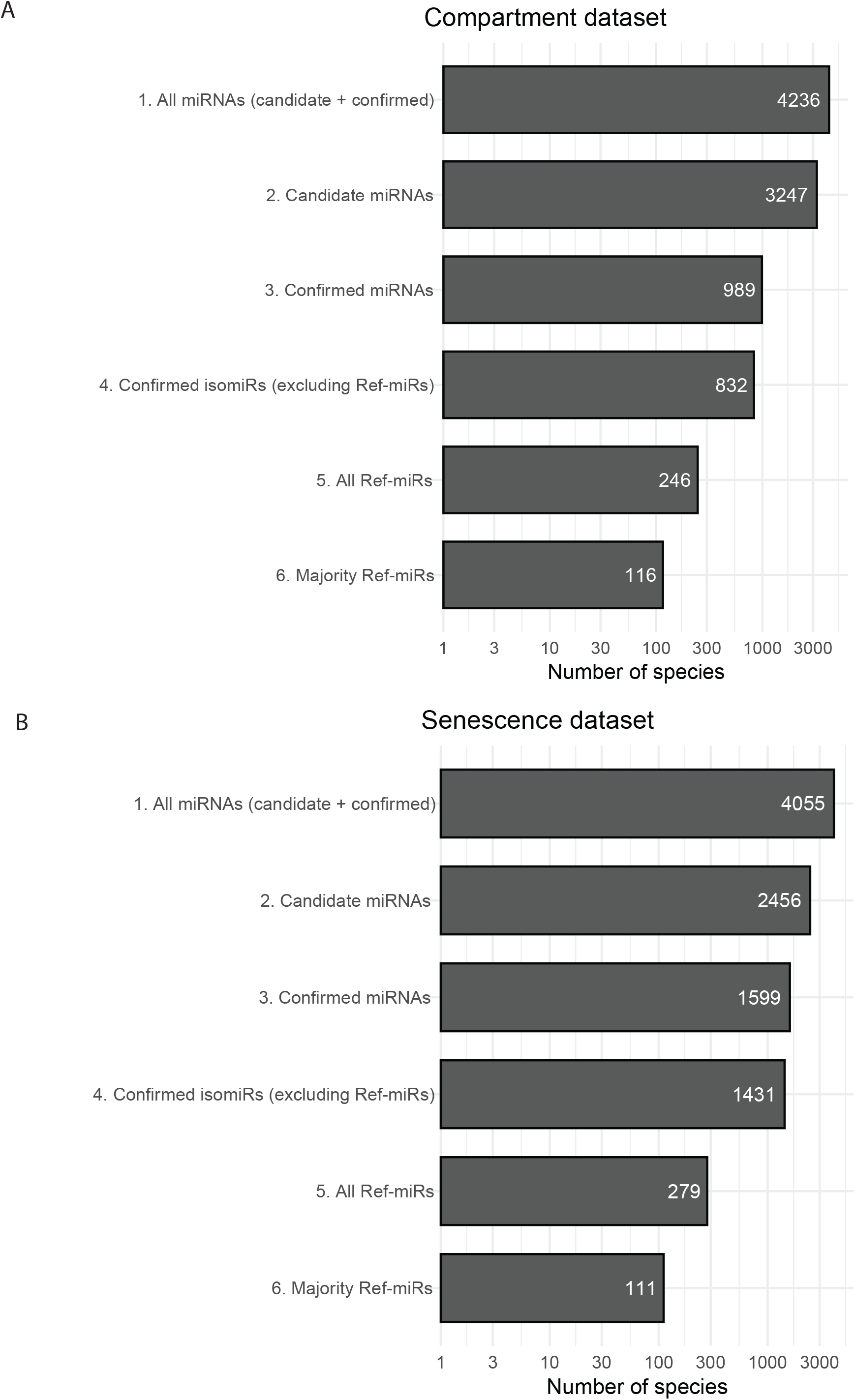
Number of detected species passing the expression cutoff. 1. All species. 2. Sequences detected by Prost! as potential miRNA species but not aligning to known human miRNA sequences (Ref-miRs) and thus discarded in analysis. 3. Sequences alignigng to human miRNA, both isomiRs and Ref-miRs. 4. All IsomiRs, excluding Ref-miRs. 5. All Ref-miRs (expressed above the filtering threshold). 6. Ref-miRs that have more than 50% of the group’s total counts. **A**. Compartment dataset. **B**. Senescence dataset.

**Supplementary figure 3:**
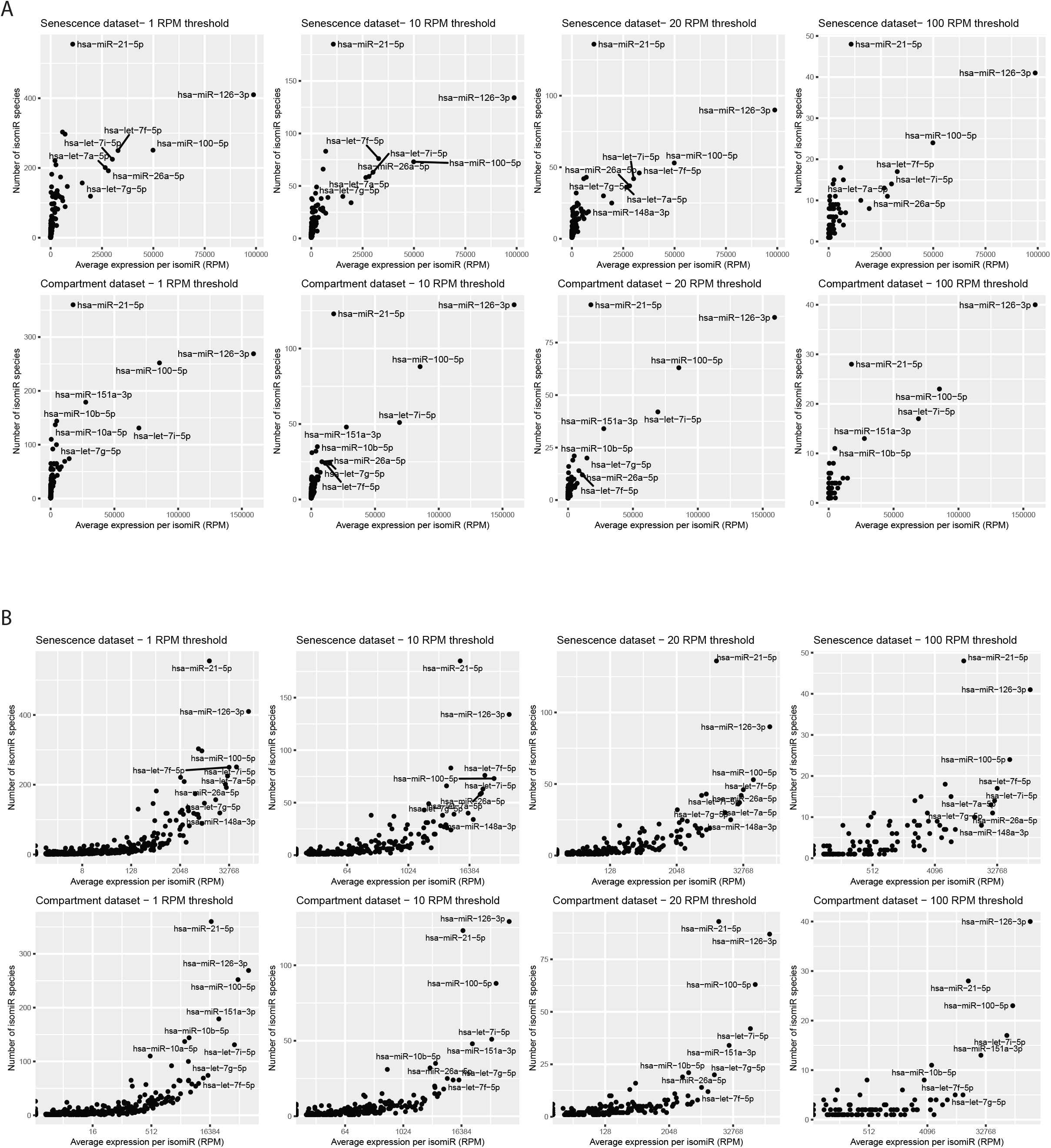
Correlation between the number of isomiR species and average isomiR expression, for each Ref-miR group, across expression thresholds, for both datasets. **A**. Regular scale **B**. Logarithmic scale for isomiR average expression.

**Supplementary Figure 4.**
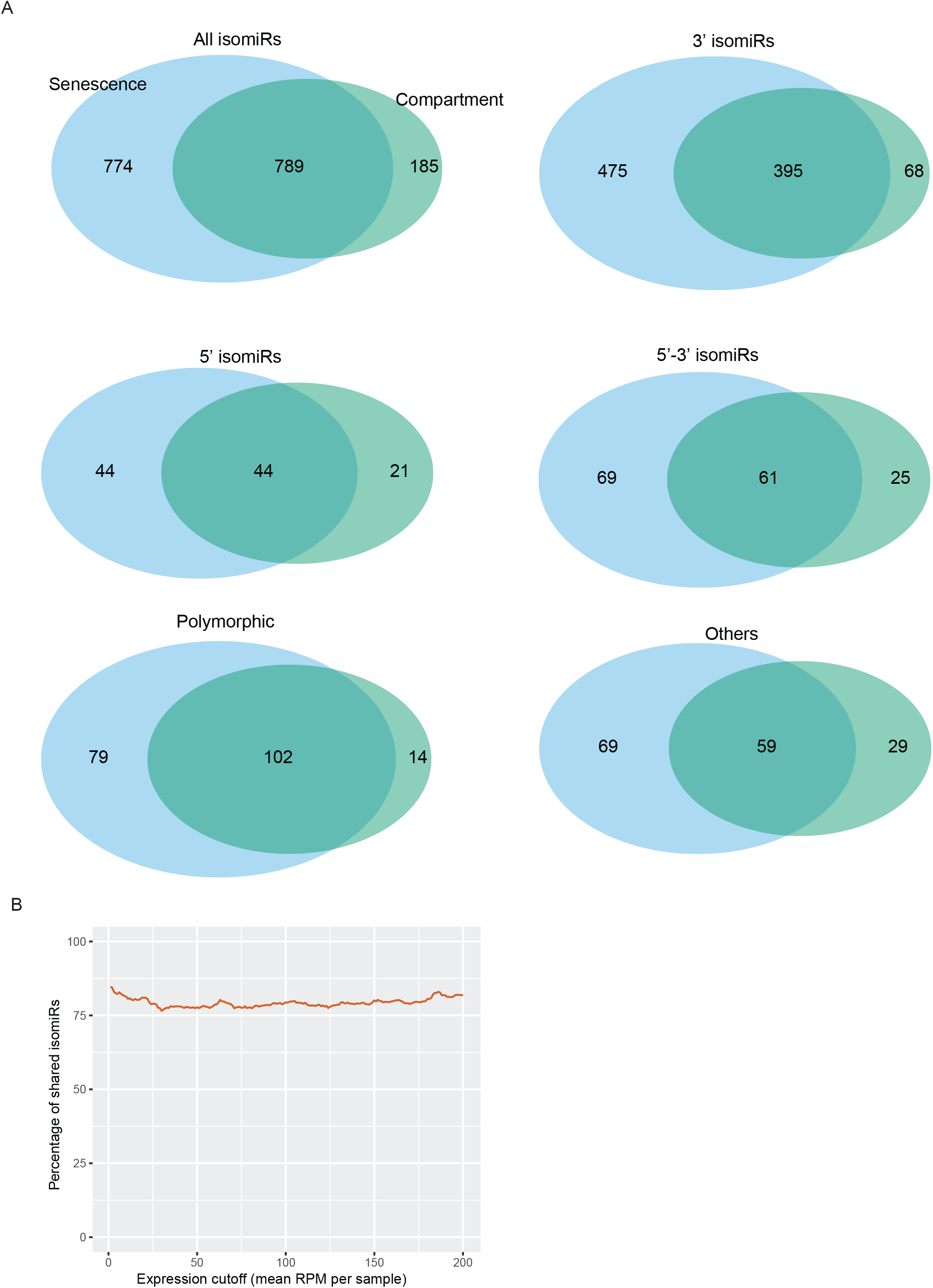
IsomiR species are shared across datasets. **A**. Counts of Senescence exclusive (left, blue), shared (middle) and compartment exclusive (right, green) isomiRs species. Each diagram is for a different species type. **B**. Percentage of shared isomiRs between both datasets, with different expression cutoffs. The percentage is calculated using *((100 * n-shared) / n-compartment)*.

**Supplementary Figure 5.**
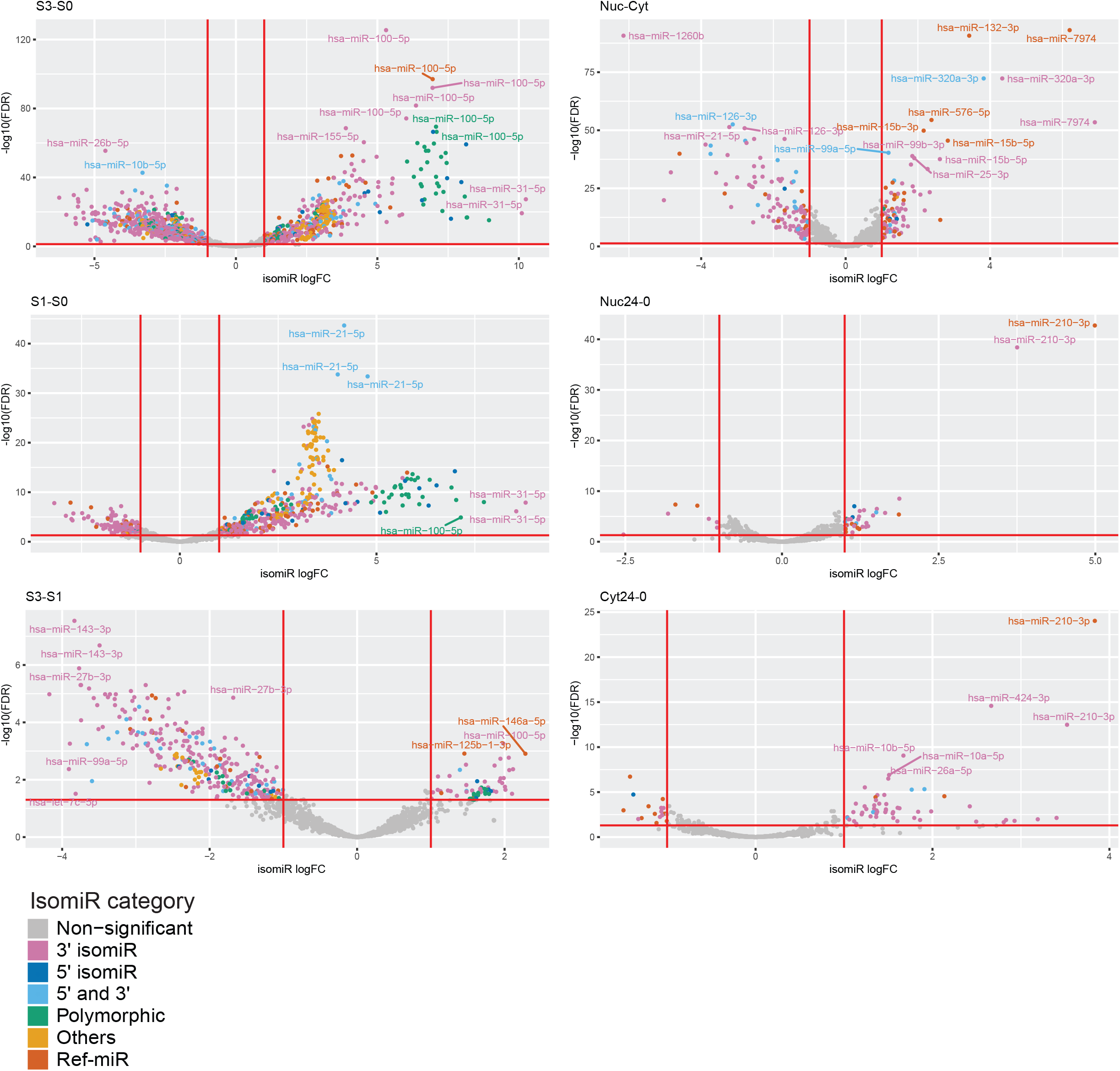
isomiR DE results for all comparisons. Species are colored by type.

**Supplementary Figure 6:**
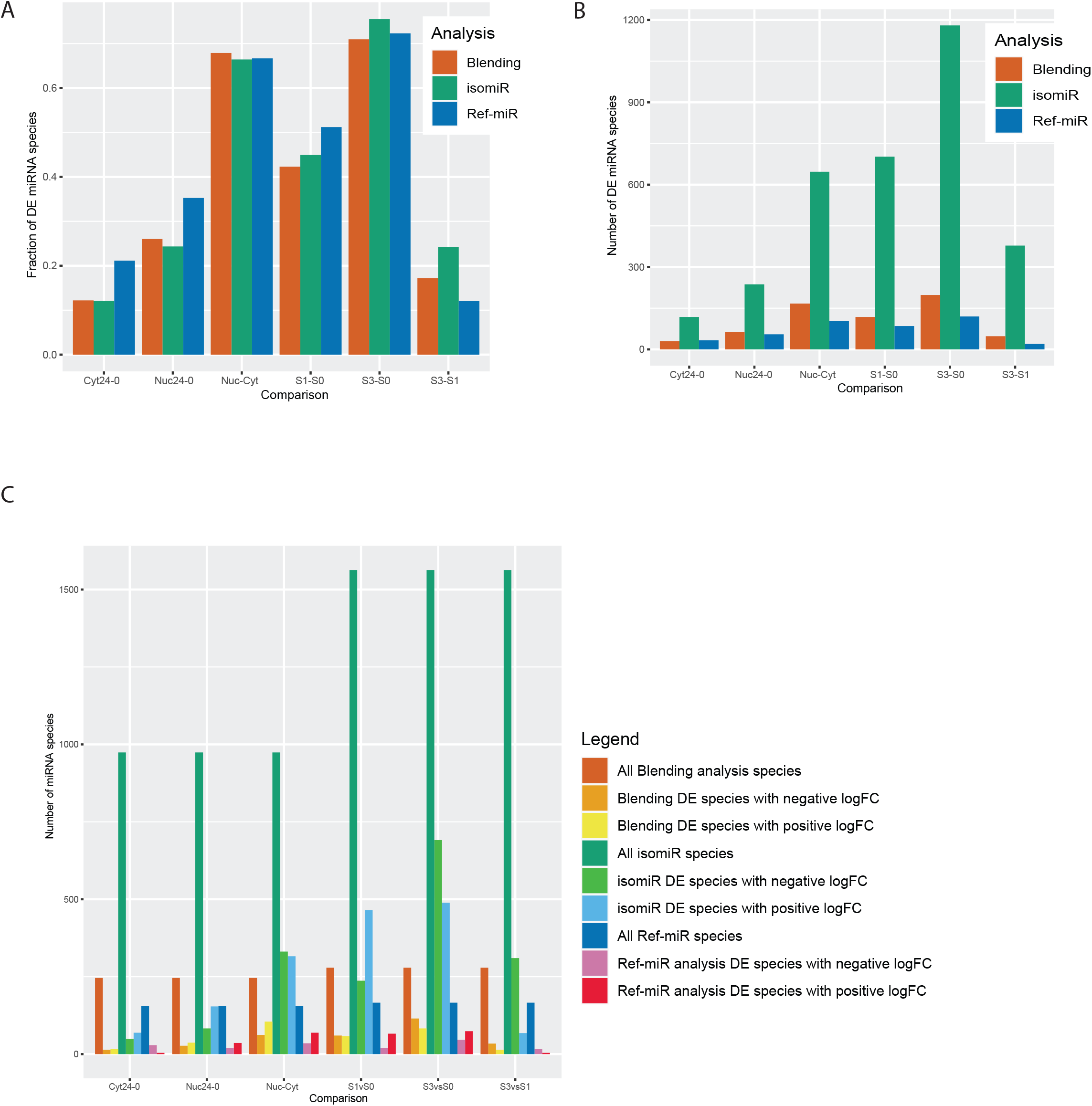
isomiR DE summary. **A**. Fraction of species that are DE across comparison and analysis methods. Blending analysis in red, isomiR in green and Ref-miR in blue. **B**. Number of DE species across comparison and analysis methods. **C**. Breakdown of isomiR DE results across comparisons. The histogram counts species that are upregulated (up), down regulated (down) and includes the total number of miRNA species (all, includes DE and non DE species).

**Supplementary figure 7:**
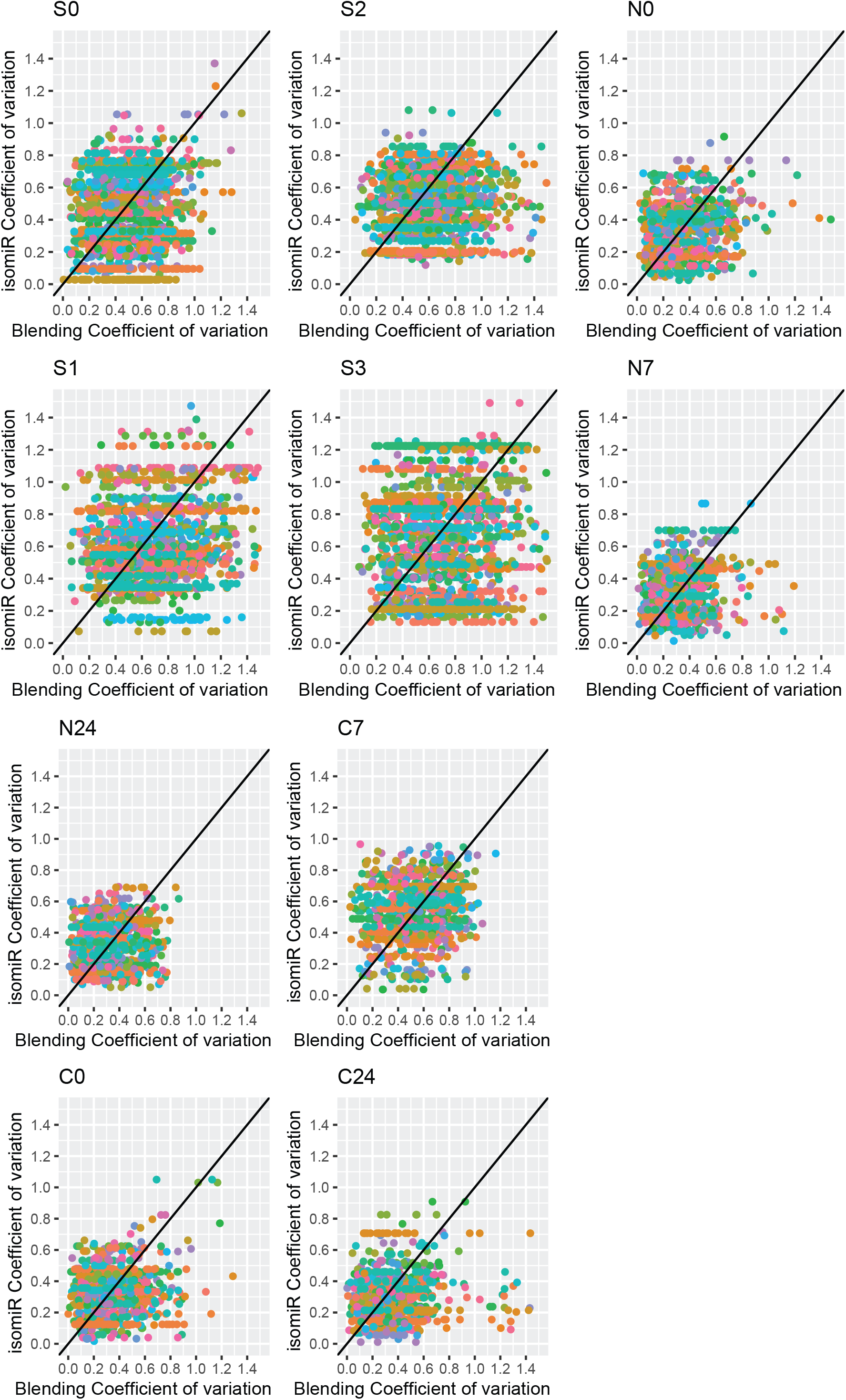
Coefficient of variation comparison between isomiR analysis and Blending analysis.

**Supplementary Figure 8:**
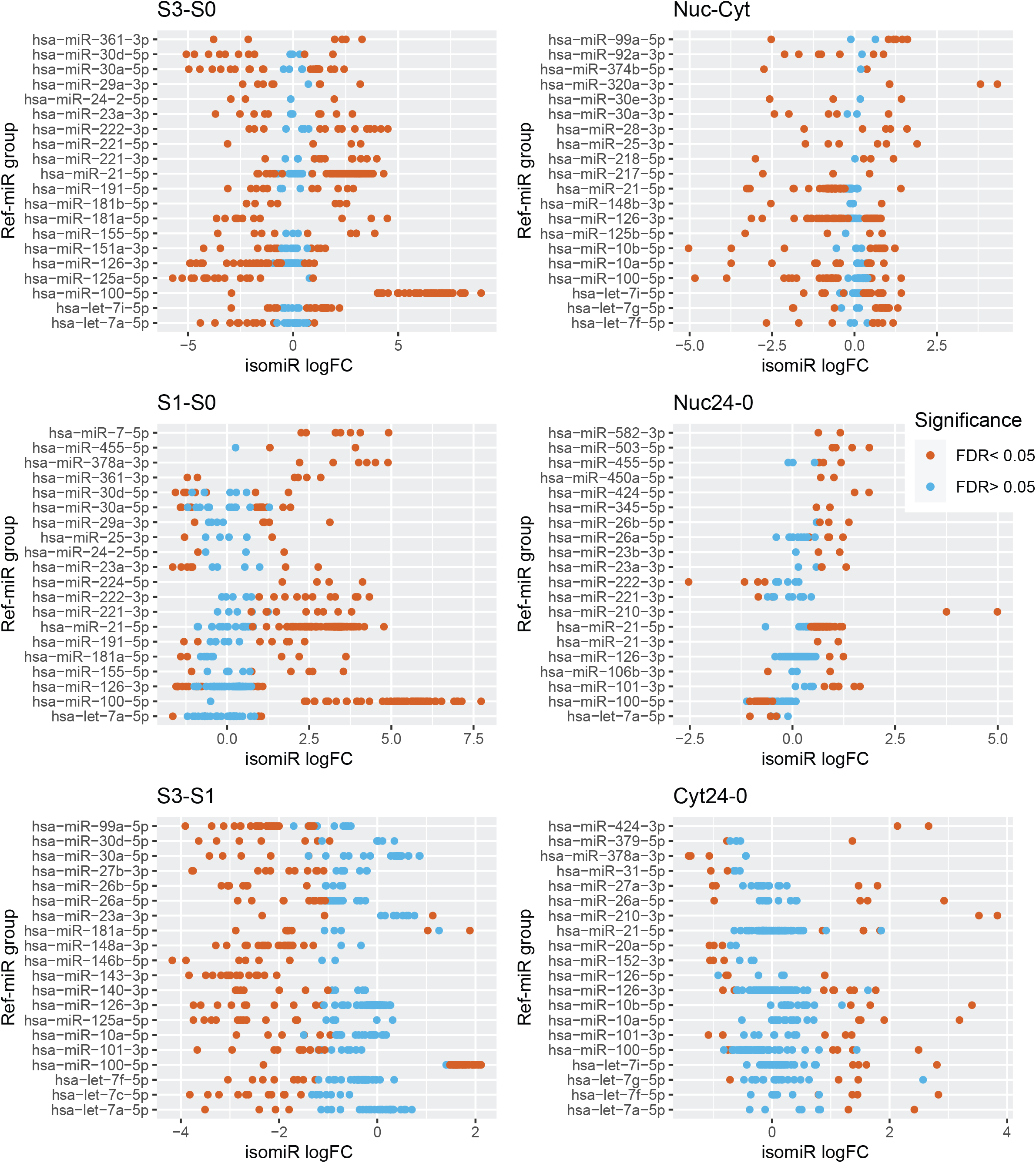
LogFC distribution snapshot, for the top 20 Ref-miR groups that contains isomiRs with the highest variation in their logFC. Each dot is an isomiR from the line’s Ref-miR group, colored by significance.

**Supplementary Figure 9:**
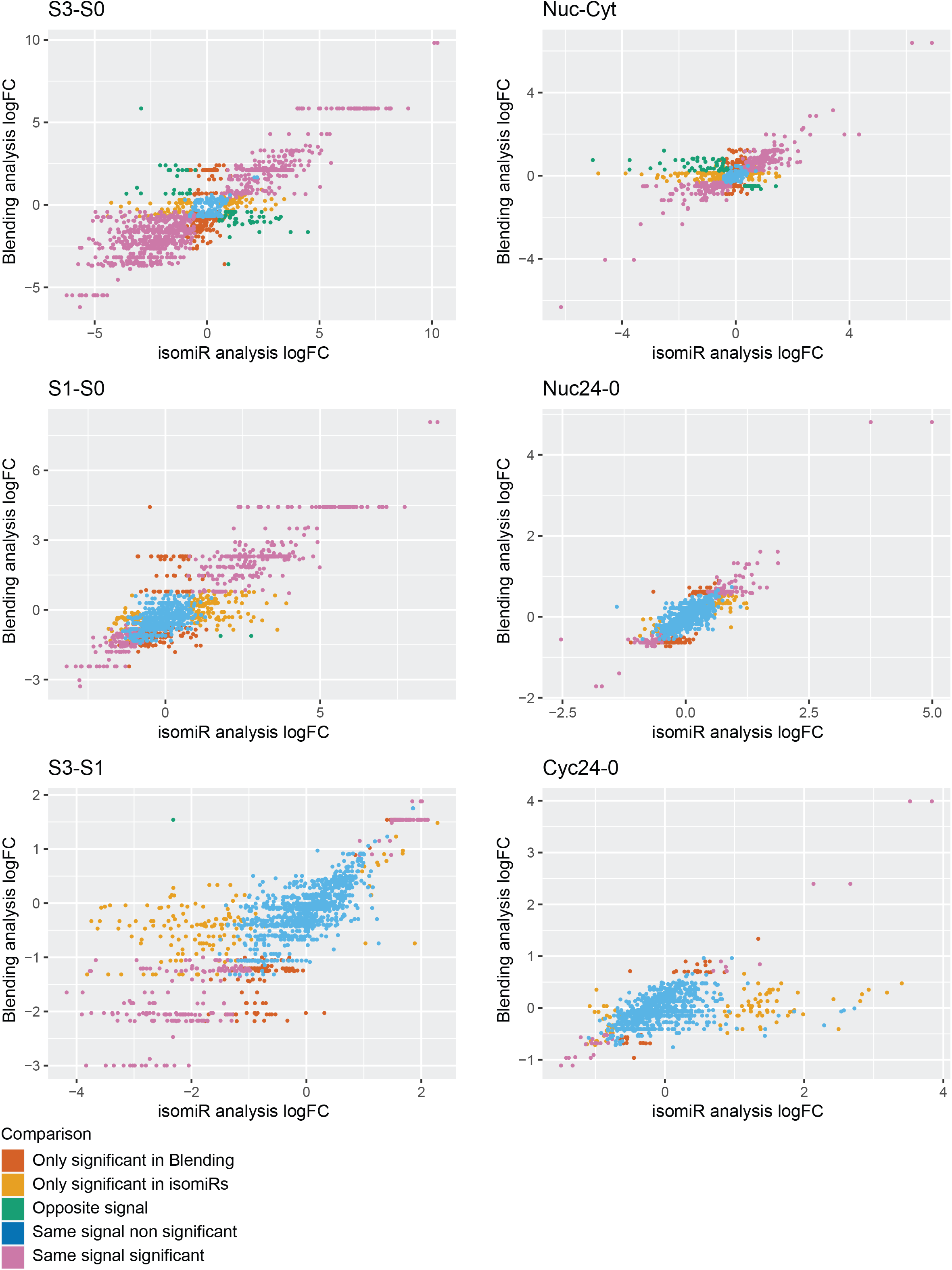
LogFC comparison between isomiR and Blending analysis, for each comparison. Each dot is an isomiR, colored by comparison group: *Only significant in Blending* means that the isomiR is not significantly DE but its Ref-miR group is DE in Blending analysis. *Only significant in isomiR* are for species for which the Blending analysis yields no significant signal, but that are DE in isomiR analysis. When both the isomiR and its Ref-miR are DE but with opposite logFC, the situation is classified as *Opposite signal*. Finally, the last cases are non DE in both methods (*Same signal non significant*) or DE in both, with the same direction (*Same signal significant*).

**Supplementary Figure 10:**
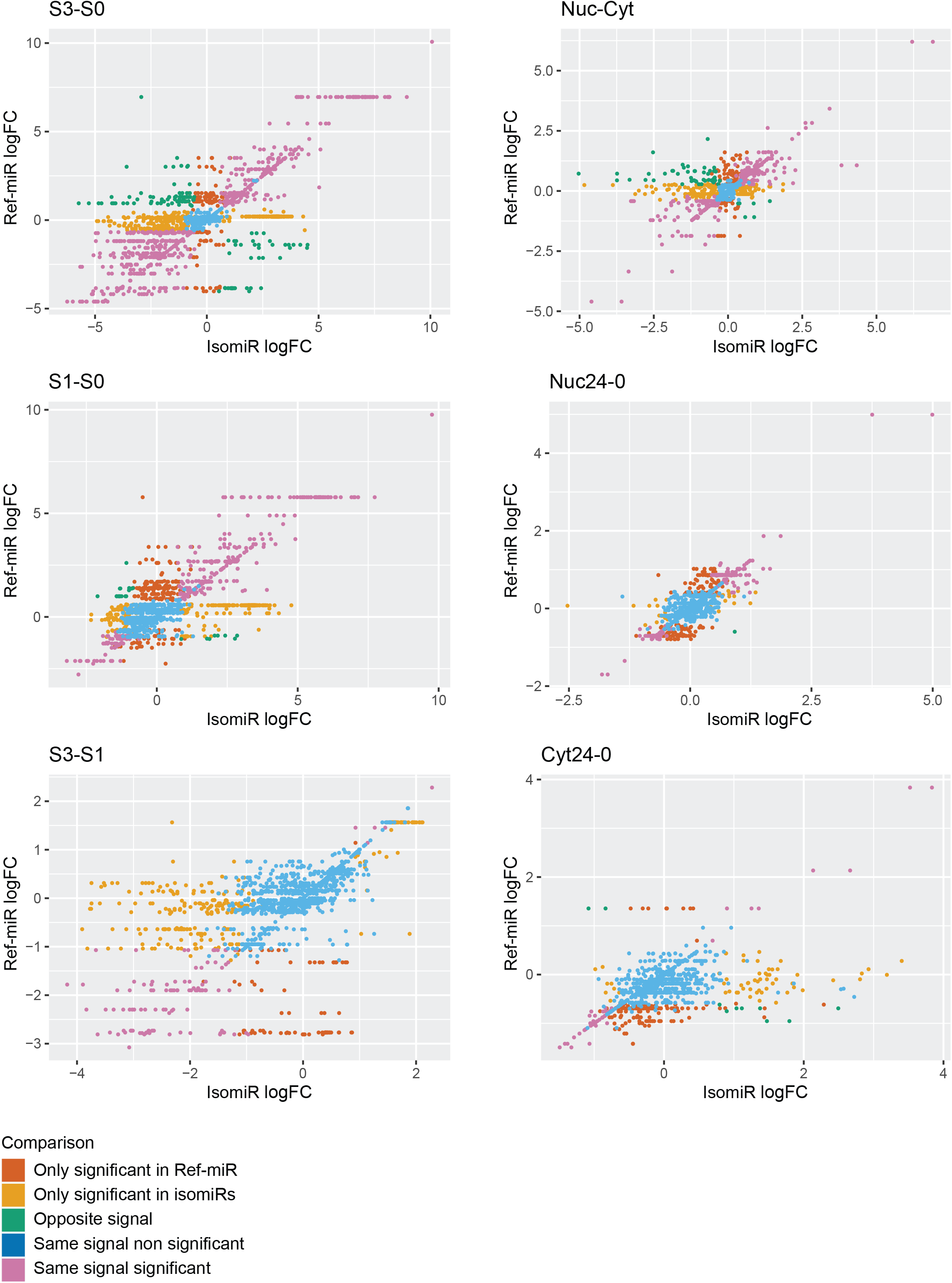
LogFC comparison between IsomiR analysis and Ref-miR analysis. Each dot is an isomiR, colored by comparison group: *Only significant in Ref-miR* means that the isomiR species is not significantly DE but its Ref-miR bin is DE in Ref-miR analysis. *Only significant in isomiRs* are for species for which Ref-miR analysis yields no significant signal, but that are DE in IsomiR analysis. When both the isomiR and its Ref-miR are DE but with opposite logFC, the situation is classified as *opposite signal*. Finally, the last cases are non DE in both methods (*same signal non significant*) or DE in both, with the same direction (*same signal significant*).

**Supplementary Figure 11:**
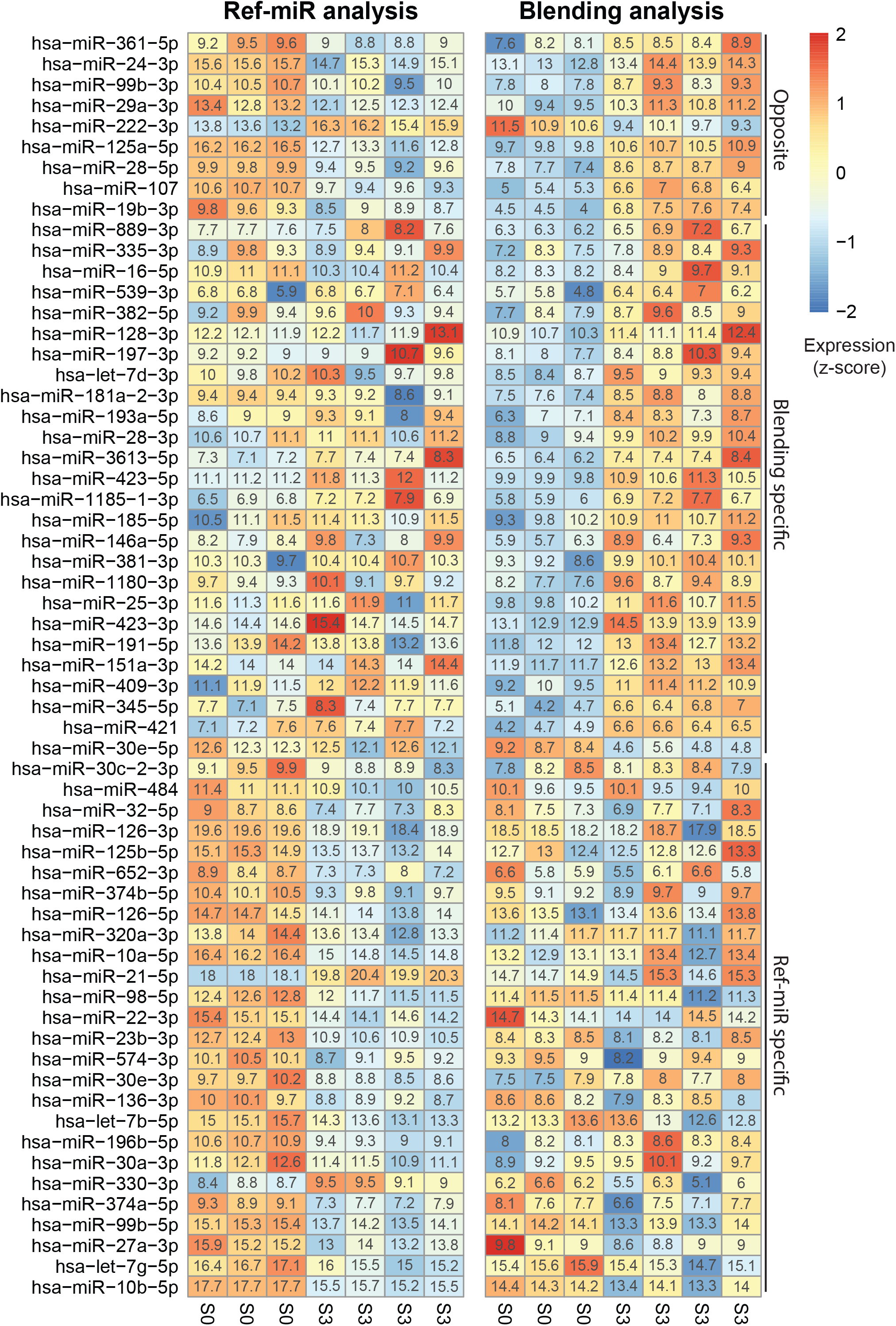
Heatmap of Deseq Normalized expression in the Senescence dataset. (condition S0 and S3). miRNA species of interest are shown, comparing Blending analysis and Ref-miR results for the same miRNAs. miRNAs are separated in *Opposite signal, Ref-miR specific* (only DE in Ref-miR analysis) and *Blending specific* (Only DE in Blending analysis).

**Supplementary Figure 12:**
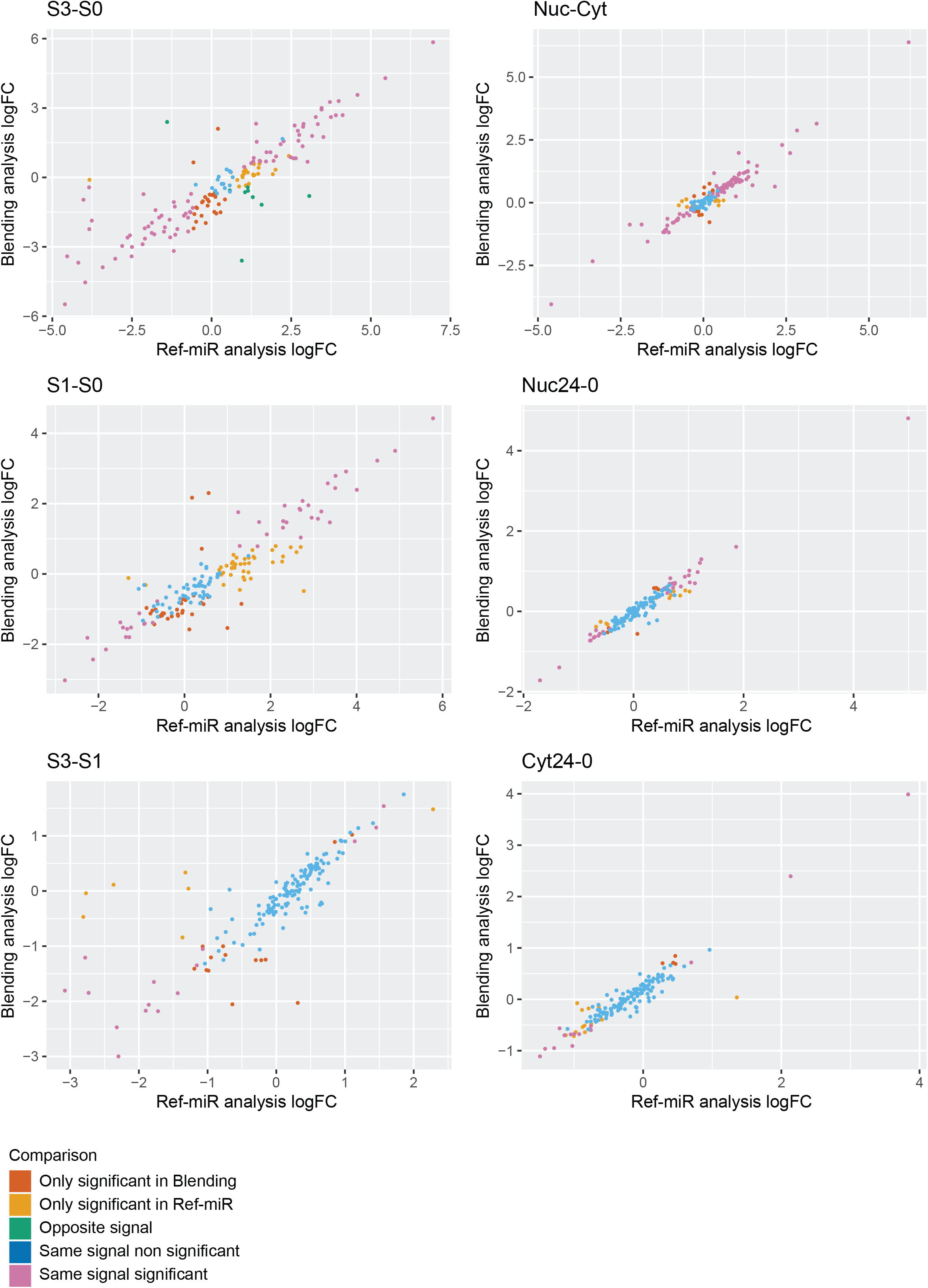
LogFC comparison between Blending analysis and Ref-miR analysis, for each comparison. Each dot is an Ref-miR, colored by comparison group: *Only significant in Blending* means that the Ref-miR species is not significantly DE in Ref-miR analysis but is in Blending analysis. *Only significant in Ref-miR* are for species for which Blending analysis yields no significant signal, but that are DE in Ref-miR analysis. When the species is DE in both analysis but with opposite logFC, the situation is classified as *opposite signal*. Finally, the last cases are non DE in both methods (*same signal non significant*) or DE in both, with the same direction (*same signal significant*).

**Supplementary Figure 13:**
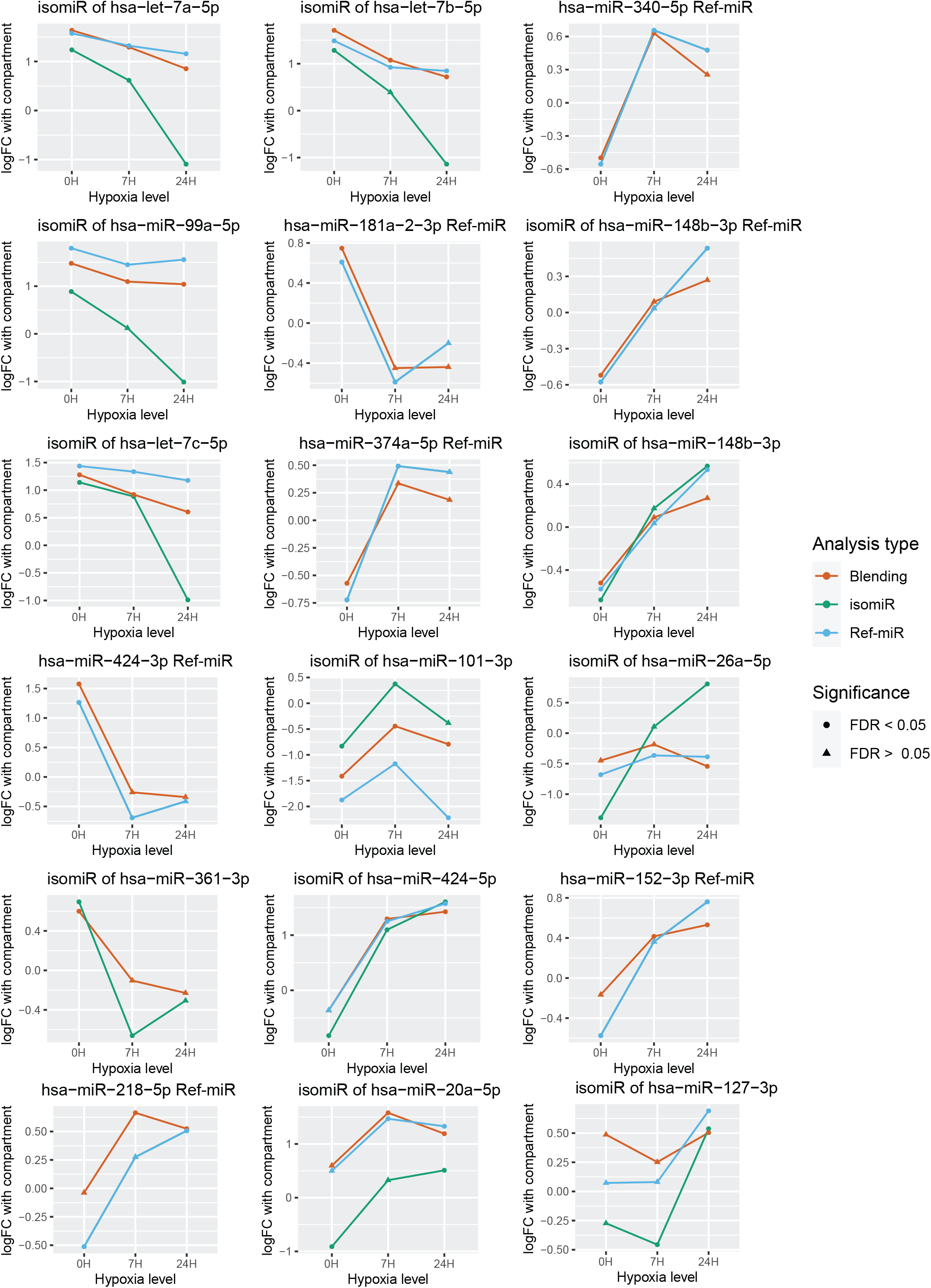
Compartment enrichement of isomiRs and their Ref-miR, between hypoxia levels. Colors correspond to the analysis type (orange is Blending analysis, green isomiR and blue Ref-miR). Dots are for significant DE and triangles for non significant.

**Supplementary Figure 14:**
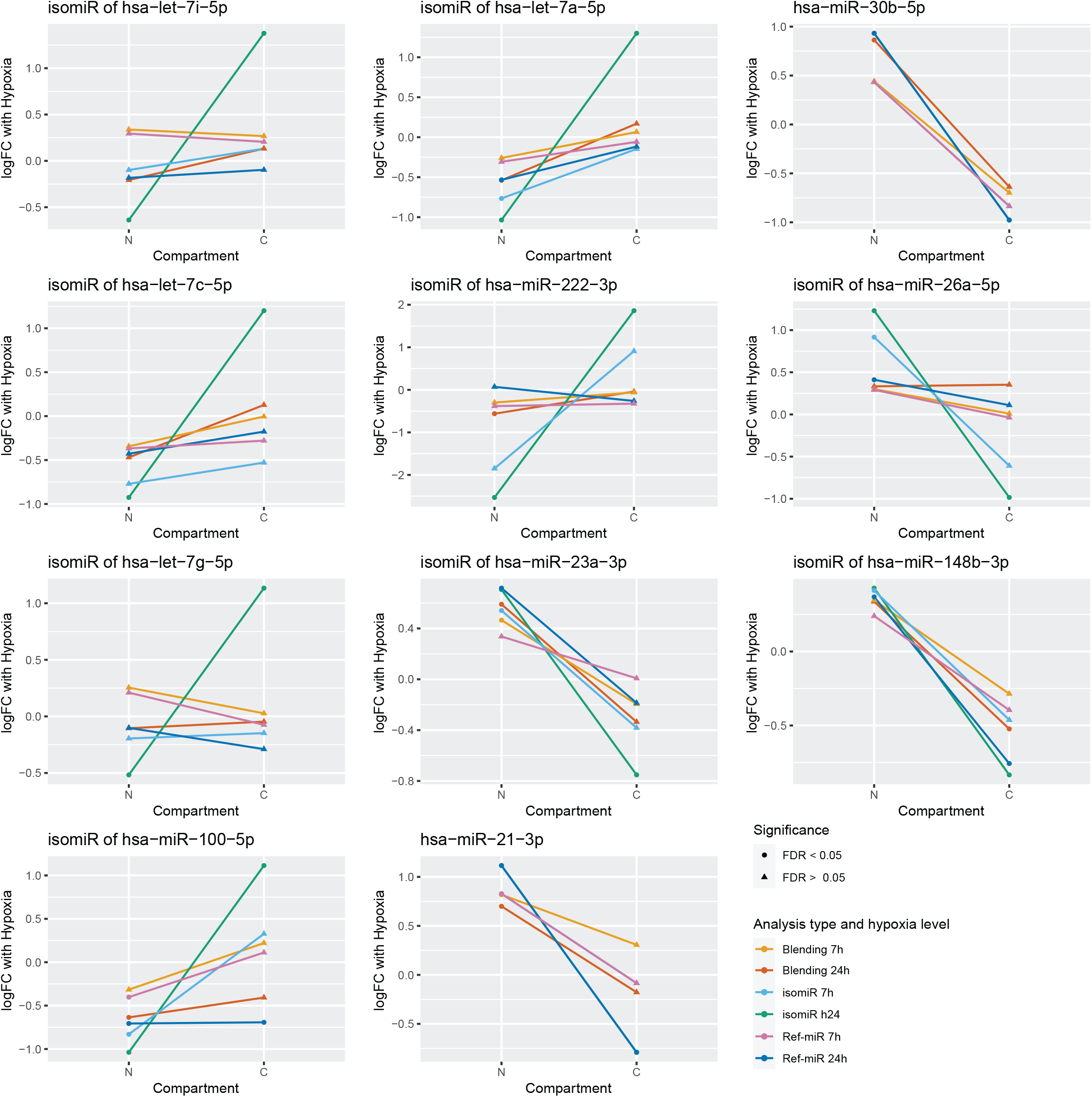
Hypoxia DE of isomiRs and their Ref-miR, between compartments. Colors correspond to analysis type and hypoxia levels. Dots are used for significant DE and triangles for non significant.

**Supplementary Figure 15:**
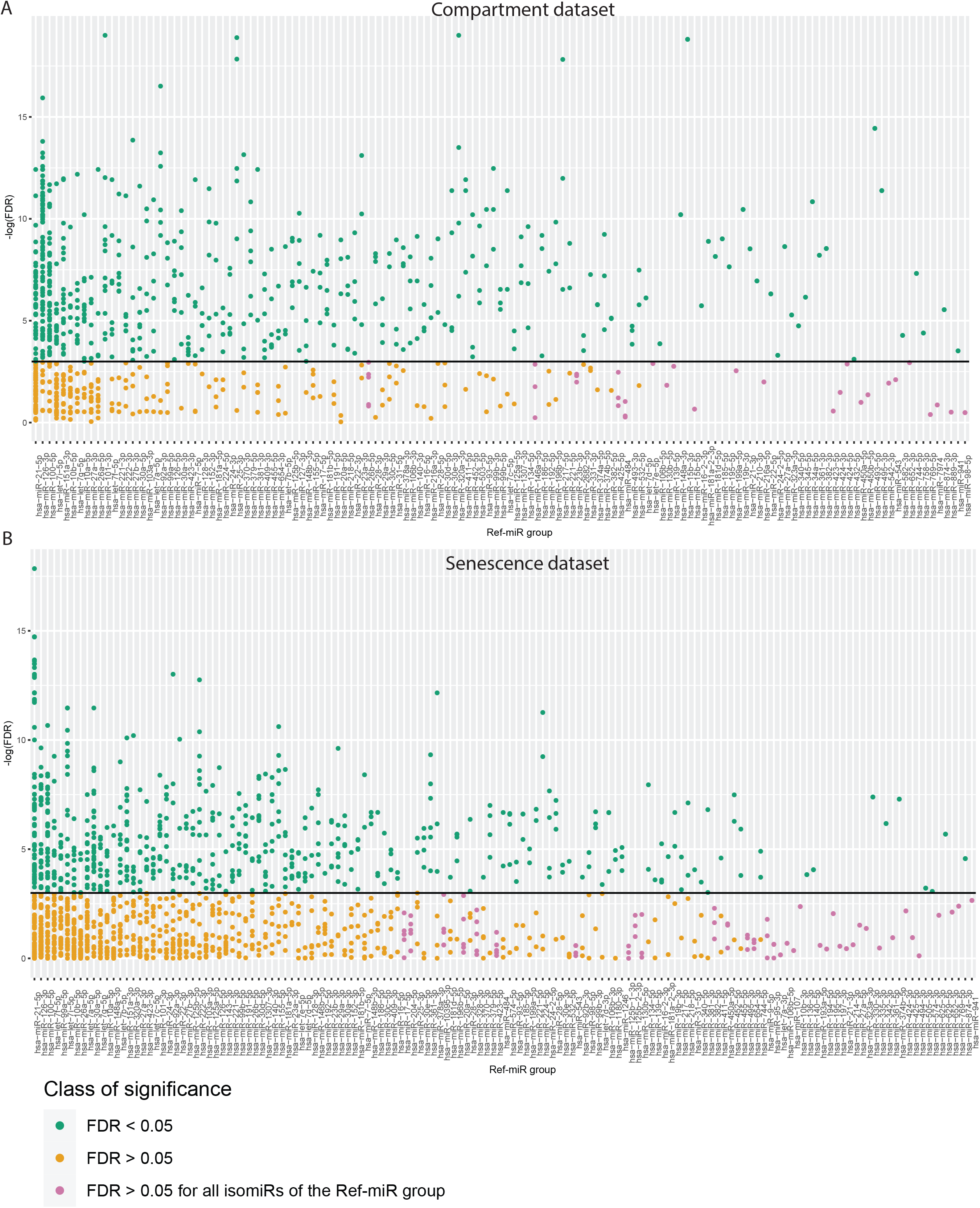
isomiR differential distribution analysis breakdown. For each Ref-miR bin, we show the statistical significance level of its isomiRs differential distribution results. Species that are significantly DD are shown in green, the ones that are not DD in orange and species for which no isomiR in the Ref-miR group is significantly DD are shown in purple. **A**. Compartment dataset, **B**. Senescence dataset (same figure than 6C but with all the Ref-miR group labels).

**Supplementary figure 16:**
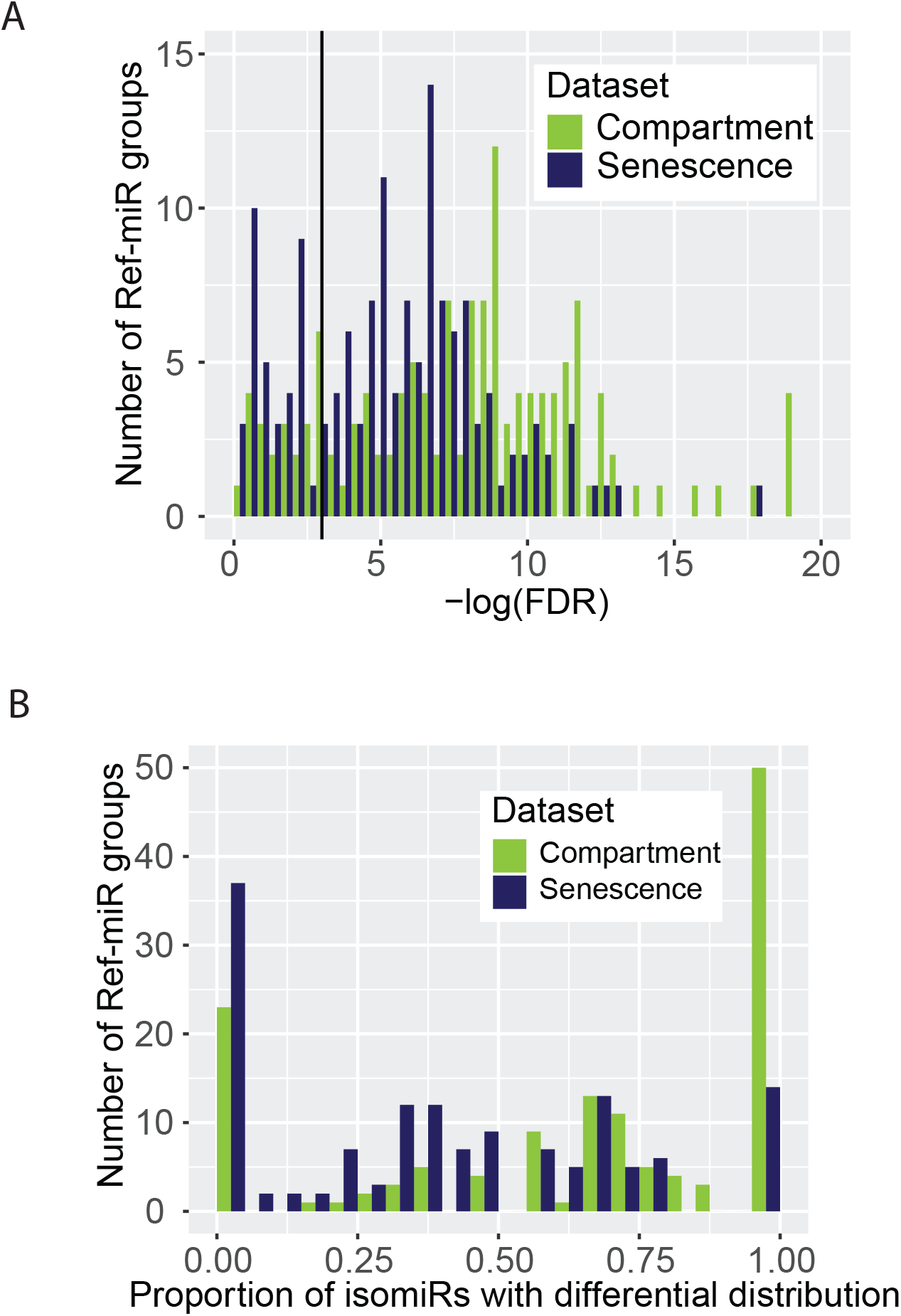
isomiR differential distribution analysis results amongst Ref-miR groups. **A**. Distribution of proportion-based Anova p-values among the most significantly DD species in each Ref-miR (one isomiR per Ref-miR), showing how differentially distributed isomiRs are dispersed among Ref-miRs bins. **B**. Proportion of differentially distributed isomiRs within Ref-miR groups.

**Supplementary Figure 17:**
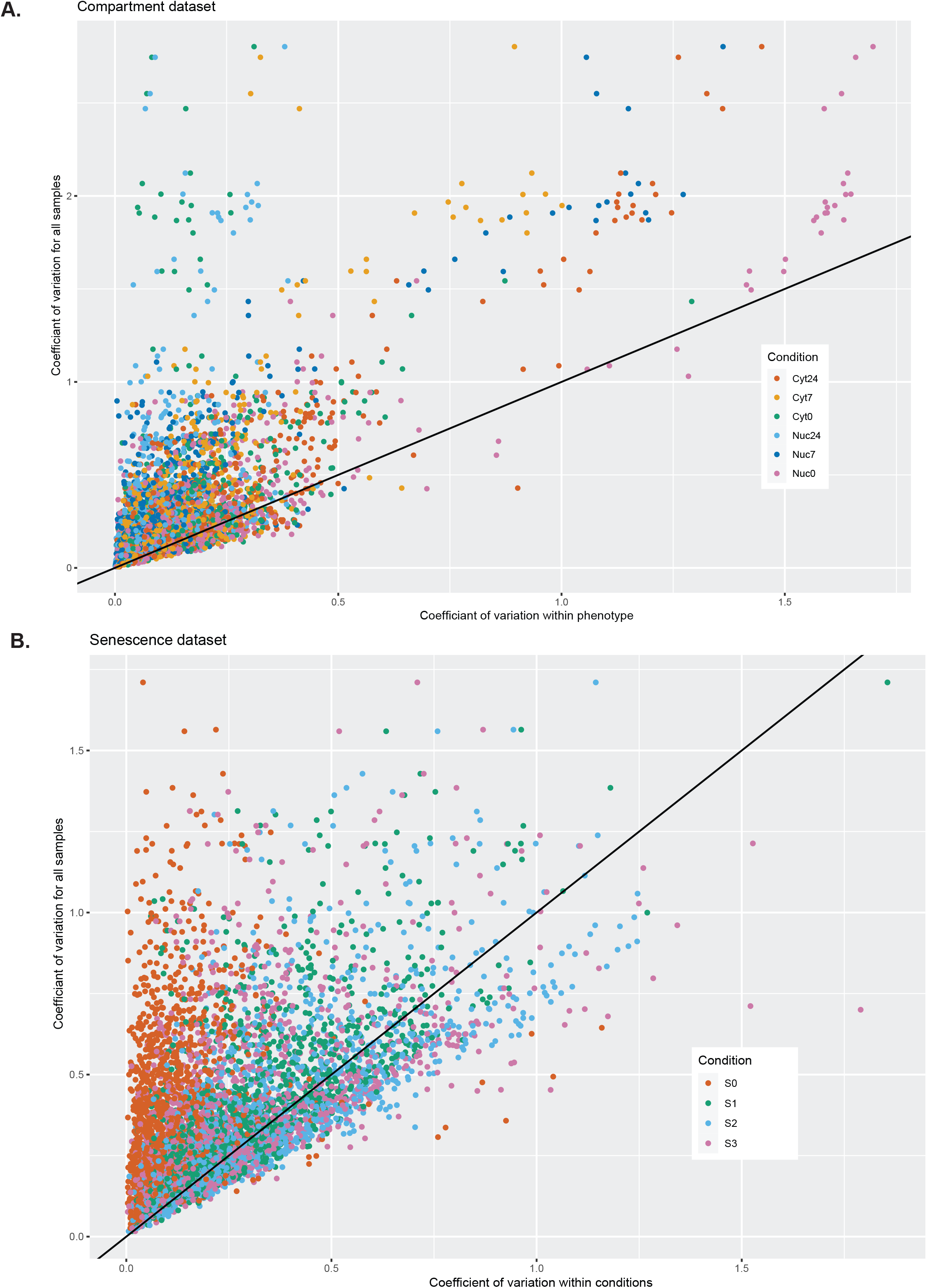
Phenotype specific variation. Scatter plot comparing the isomiR proportion coefficient of variation across all samples and within each condition-specific group. For **A**. the compartment dataset and **B**. the senescence dataset.

